# Minimally verbal children with autism may see the global, but point local: A behavioral and eye-tracking study in visual perceptual processing

**DOI:** 10.64898/2026.01.14.698823

**Authors:** Hannah S Sykes-Haas, Oren Kadosh, Yoram S Bonneh

## Abstract

Recent work suggests that pointing based assessments may underestimate or mischaracterize perceptual and cognitive abilities in minimally verbal autism (mvASD). Extending this line of work, we address the classic global local question by testing whether mvASD children access global visual structure and whether it is consistently expressed in behavior using Kanizsa illusory contours (KIC) and circular colinear contours (CC). Typically developing (TD) and mvASD children viewed KIC and CC stimuli on a touchscreen while spontaneous eye gaze and pointing were recorded. Participants also completed a goal directed drag and drop matching task that required selecting the solid shape corresponding to the global KIC configuration. In spontaneous responses, mvASD children were more drawn to local elements (e.g., individual inducers, the CC frame, single Gabors) and showed less centralized responding than TD peers. Notably, CC pointing was often anchored to one half of the contour regardless of contour screen location, suggesting object centered global selection. Also, in the drag and drop task, approx. 90% of mvASD participants performed above chance in at least one condition and half performed at ceiling, demonstrating accurate global matching. These findings suggest that global structure is available in mvASD but may be underweighted in spontaneous response selection under ambiguity, consistent with the Affordance Competition Hypothesis and ambiguity resolution accounts.

**Lay Summary:** Autism research often asks whether autistic people can see the wood for the trees—integrate parts into a coherent whole. This is difficult to test in minimally verbal autism because many standard tasks rely on a single pointing response that may not reflect what a child actually perceives. In our study, children viewed visual illusions in which a clear shape seems to appear even though it is not actually drawn, for example, a triangle you *see* because several Pac Man shapes are arranged in just the right way, while we measured both where they looked and where they touched the screen. Minimally verbal autistic children were often more drawn to the smaller parts than typically developing peers, yet when a simple drag and drop game required an explicit choice of the overall shape, most selected the correct global shape. Together, the findings suggest that global structure can be available in mvASD, but its expression in spontaneous behavior may be more context dependent under uncertainty, highlighting the value of using more than one way to measure ability.

## Introduction

The long standing local global perception debate (Van der Hallen et al., 2015) has excluded the performance of approximately 30% of the autism population, namely the minimally verbal with profound autism (Tager-Flusberg and Kasari, 2013). This general underrepresentation in autism research, including the study of visual perceptual processing is largely due to the experimental challenges involved in testing this significant clinical subgroup. Often tasks require pointing responses, compliance with instructions, or verbal reporting, known to be challenging for individuals with mvASD (Courchesne et al., 2015, Tager-Flusberg, 1999 and Tager-Flusberg et al, 2017). In the last decade or so there has been an upsurge in mvASD inclusion in research and Courchesne and colleagues have successfully assessed mvASD children using what they coin *strength-informed* visual processing paradigms (Courchesne et al., 2015 and 2019, Dawson and Mottron, 2011). These paradigms, including the Children’s Embedded Figures Test, visual search tasks, and Raven’s Colored Progressive Matrices, employed nonverbal visual materials presented in printed and cardboard formats, allowing children to indicate responses through physical placement. Using these approaches, a substantial proportion of mvASD children completed the tasks successfully, in some cases performing comparably to typically developing peers. While these findings have been interpreted as reflecting alignment with visual–perceptual strengths in autism (Bertone et al., 2005, Mottron et al., 2006 and Dawson and Mottron, 2011), the *tangible* response formats employed represents an additional notable methodological characteristic of these particular assessments, the potential contribution of which is considered further in the Discussion.

In our recent studies on visual perception and use of eye-tracking with the mvASD population we have further developed alternative response contexts and modelled practice trials to reduce the need for receptive language skills. We have addressed maintenance of task engagement by applying very brief trial runs (0.5–2.0 min) and unlimited response time, as well as unrestrained head movement during eye-tracking. Our findings suggest that mvASD individuals demonstrate early and simple perceptual and cognitive competence. For example, children fixated correctly on visual oddball target, even when they failed to point to visual target (Sykes-Haas and Bonneh, 2025, preprint). We also found mvASD individuals that successfully match a written word to image via eye-tracking performed at chance when tested via pointing (Ellert et al., 2025). Furthermore, visual saliency, deviance from distractors and complexity appear to modulate touch performance (Sykes-Haas and Bonneh, 2025, preprint). During this recent study, employing oddball tasks to assess levels of visual processing in mvASD we observed that approximately half of mvASD children who *successfully* detected a global triangular Kanizsa-induced contour (KIC) oddball consistently pointed to local inducing elements rather than the center of the illusory shape, unlike typically developing children’s consistent pointing to the global center (Sykes-Haas and Bonneh, 2025). Although successful detection implies access to the target’s global configuration, concurrent local pointing challenges whether perception itself is locally organized (Mottron et al., 2006), such that responses are driven by local feature cues (e.g., Gestalt or deviance), or global perception is weaker (Booth and Happe, 2018 or Naya et al., 2021) or whether global perceptual access is present but translated into action via response-selection processes that favor local or salient. Behavioral selection may reflect differences in how visual information guides behavior and action selection in minimally verbal autism.

In the companion paper mentioned above we characterized basic and mid-level visual perception in the in mvASD (Sykes-Haas and Bonneh, 2025). The present study examines the same cohort and extends that work by investigating global–local organization and how global visual structure processing is weighted in response selection in minimally verbal autistic children. We adopt an explicitly exploratory approach . Given the paucity of empirical data in this population, we do not advance directional hypotheses regarding local–global integration in mvASD. We examined mvASD children’s response selection and processing of global visual structure using two contour-based paradigms: Kanizsa illusory contours (Kanizsa, 1979, KIC) and circular colinear contours (CC) constructed from Gabor elements (Nayar et al., 2017, Nieder, 2002, Guttman & Kellman, 2004, Gowen et al., 2020, Kovács & Julesz, 1993). Across these paradigms, we measured spontaneous eye-gaze and pointing behavior, as well as performance in an alternative behavioral response context—a drag-and-drop matching task requiring explicit selection of solid shapes to corresponding global KIC shape alternatives. By contrasting spontaneous and explicit response contexts, this design allows examination of whether perceptual access to global structure is present and may diverge from the way visual information is expressed behaviorally under conditions of perceptual ambiguity in mvASD vs TD (Robertson and Baron-Cohen, 2017). Together, this multi-method approach was designed to assess whether access to global visual structure in mvASD can dissociate from overt action selection across spontaneous and goal-directed contexts.

## Methods

### Participants

Twenty-four minimally verbal autistic children (7 females and 17 males) were enrolled (see Sykes-Haas & Bonneh, 2025). Some participants were excluded post-enrolment due to inability to complete the experimental procedure or insufficient usable data. Twenty-one contributed data to at least one task in the finale analyses, and Ns varied by task due to protocol-related exclusions (Ns reported per analysis), reflecting the practical challenges of assessing this subgroup. At the time of the Kanizsa-induced contours, pointing and eye-gaze, and drag-and-drop game tasks, children mean age was 9.6 years (SD = 1.5; range = 7.5-12; n = 21). The original circular contour task (CC) data collection, conducted two years previously, was affected by technical problems; we therefore re-administered the CC task, and children were correspondingly older (M = 11.3 years, SD = 1.9; range = 7.5–14; n = 18). All had confirmed clinical ASD diagnoses according to DSM-IV-TR or DSM-5 criteria, independently verified by two accredited diagnosticians (one physician and one psychologist), as required by the Israeli Ministries of Health and Education. Minimal verbal status was defined as spontaneous use of fewer than 30 distinct communicative words or phrases and the absence of consistent phrase-level speech, as reported by parents and teachers (Tager-Flusberg & Kasari, 2013). Participants that were at least 8 years old and met the Lancet Commission “profound autism” administrative classification via the minimal-language criterion (persistent absence of consistent phrase speech after age 8), based on parent/teacher report (Lord et al., 2022).

Thirty-two typically developing (TD) children (16 females and 16 males) served as age-matched controls. During KIC tasks, mean age was 9.1 years (range: 5.5-12.7y and SD: 2y, N=25) and during CC task, mean age was 9.75 (range: 6.40 -13.5 ands SD: 2y, N=20). All attended the regular school system. None had known developmental, neurological, or visual atypicalities or diagnoses; all had normal or corrected-to-normal vision.

We collected scores for non-verbal fluid reasoning using Raven’s Colored Progressive Matrices (RCPM; Raven et al., 1998) in both pointing and puzzle-board formats, communication abilities using the Low-Verbal Investigatory Screener 4.0 (<u>L-VIS</u> 4.0; Naples et al., 2022), and confirmed autism diagnosis using the Social Communication Questionnaire (SCQ; Rutter et al., 2003). These measures were not collected in TD participants, who served solely as a comparative performance group and behaved uniformly—performing at ceiling on the circular contour and drag-and-drop tasks and consistently pointing to the center of CC and KIC—rendering additional descriptive measures uninformative.

All mvASD participants were assigned pseudonyms beginning with the initials “H” or “L” (high and low performers, respectively), based on their overall performance on visual oddball tasks described in Sykes-Haas & Bonneh (2025). As shown in the Results section, mvASD participants continued to fall into these performance groups in the current study’s experimental tasks as well (see Circular Contour Detection and Kanizsa Drag-and-Drop Game).

### Exclusion criteria for mvASD

The mvASD participant group were not included if the child was diagnosed with comorbid neurological or genetic conditions except for medically controlled epilepsy, developmental delay and intellectual disability, as standardized tests and/or diagnostician’s clinical impression may at times underestimate cognitive abilities in mvASD (Courchesne et al., 2015 and 2019, Bauminger - Zviely et al. 2020, Pizzano et al 2024, Sykes-Haas & Bonneh, 2025). Sixteen mvASD participants were treated daily with psychotropic medication (Risperidone, Ritalin, Carbamazepine, Atent, melatonin, or medical cannabis). Ethical approval was granted by the Chief Scientist of the Israeli Ministry of Education and the Ethics Committee at Bar-Ilan University. Written informed consent was obtained from all parents (both mvASD and TD groups). Data collection was conducted either in the child’s home or at their educational institution, depending on parental preference. All TD participants and one mvASD participant were assessed at home.

**Table 1.** Summary of descriptive measures^1^.

| **Descriptive measures for mvASD**<br>(KIC, n=20 and CC, n=18)) | **Mean (± SD)**<br>(excluding f.p.<sup>2</sup>.) | **Range** | **Estimated IQ** |
| --- | --- | --- | --- |
| **Boys/Girls** | 16/6 | ---- |  |
| **Chronological decimal age** | KIC: 9.6 (1.5)<br>CC: 11.3 (1.9) | 7.5-12<br>7.5-14 |  |
| **L-VIS (communicative competence)<sup>3</sup>** | 8.7 (3.1) | 8-14 |  |
| **RCPMpuzzle<sup>4</sup>-**<br>nonverbal fluid reasoning-percentiles | 52 ± 34 (n=10) | f.p.<sup>6</sup>.-97<sup>th</sup> | f.p.-129 |
| **RCPMBooklet<sup>4</sup>-**<br>nonverbal fluid reasoning- percentiles | 63 ± 33 (n=6) | f.p.-97<sup>th</sup> | f.p.-128 |
| **Perseveration on the RCPMpuzzle**<br>(same 2 consecutive answers) | 6.8 ± 6.5 | 0-20 | |
| **SCQ score<sup>5</sup>** | 23.9 (4.2) | 15-30 |  |

<sup>1</sup> RCPM data not available for some mvASD participants (see Appendix 1 for more details) <sup>2</sup> f.p.=floor performance
<sup>3</sup> Maximum score on L-VIS communicative abilities is 17. <sup>4</sup> excluding f.p (less than 1<sup>st</sup> percentile) and only scores within age-range (above 5<sup>th</sup> percentile. No mvASD scored between 1<sup>st</sup> and 5<sup>th</sup>). RCPM Score from testing in 2024 (Sykes-Haas and Bonne, 2025)
<sup>5</sup> Cut-off score for increased likelihood of autism is $\geq 15$ (Rutter et al., 2003). <sup>6</sup> All participants with a RCPMpuzzle f.p. made at least 3 mistakes by perseveration (see appendix 1).

### Apparatus

Stimuli were presented on a 14-inch Lenovo Yoga C940 touchscreen laptop (1920 × 1080 pixels, 60 Hz) using the PSY experimental tool for psychophysics and eye-tracking developed by author YSB, as in our previous studies (e.g., Bonneh et al., 2016; Meidan & Bonneh, 2025), together with custom MATLAB code implemented via PsychToolbox (Brainard, 1997; Pelli, 1997). Eye-tracking was performed using an attached Tobii 4C eye tracker (90 Hz sampling rate; manufacturer-reported accuracy of ∼0.5°–1°). This system was selected for its unobtrusive design, allowing natural head movement while providing sufficient precision for the present study, which focused on sustained gaze position rather than fine-grained saccadic measures. To ensure stable positioning, the eye tracker was mounted using a Tobii EyeMobile computer tray attached to an adjustable arm. Participants were seated at an average viewing distance of ∼53 cm (mvASD) and ∼54 cm (TD), and room illumination was maintained at approximately 500 lux. Eye-tracking was calibrated prior to each session using a 3-point calibration routine.

### Stimuli (see supplementary materials for more details)

#### Circular Colinear Contour (CC) Task

Circular colinear contour (CC) stimuli were constructed from Gabor patches arranged to form a single closed circular contour embedded within a field of randomly oriented Gabor distractors (Figure 1a; Supplementary Table S1). The contour stimuli were generated as configurations of symmetric Gabor patches (wavelength and envelope of 16 pixels) with 30% contrast on a gray (60 cd/m²) background, embedded in a random background (Field et al., 1993). First, a background of randomly oriented Gabor patches arranged on a grid with a specific spacing that varied across conditions was set to cover 1900x600 pixels rectangle, with a position jitter radius per patch of 48 pixels. Then, a circle of patches with a radius of 160 pixels, inter-patch interval of 88 pixels, and uniform tangential orientation jitter of +/-20 deg, were added in one of 6 positions (0,+/-450 and ,+/-250 pixels for x and y respectively). Background patches that were closer than 48 pixels to a target patch (center-to-center) were erased. There were 3 contour Gabor density difficulty levels in separate runs according to the background inter-patch interval: Easy (640, 192 pixels), Medium (128, 96 pixels), and Hard (64,48 pixels), two intervals mixed per level. The contour appeared pseudo-randomly in one of six fixed screen locations. Performance was assessed using spontaneous pointing responses. Participants were given unlimited response time, so each trial remained visible until a touch response was registered.

#### Kanizsa Induced Contours (KIC): Spontaneous Response Tasks

Participants were assessed on KIC during a pointing task and an eye-gaze task. In both tasks, single triangular Kanizsa figures (“Single Kanizsa Triangle,” Figure 1b and 1c) were presented pseudo-randomly at the center of the screen or in one of four screen quadrants, and in three size variants (Figure 1b; Supplementary Table S2). Four stimulus conditions were tested: Original, Central dot, Line, and Grey-Luminance Modulation. These conditions progressively increased the physical definition and perceptual salience of the contour. In the Line and Grey-Luminance conditions, the triangle was explicitly defined and therefore no longer illusory. In the eye-gaze task, the stimulus changed size and/or location every 2 s. In the pointing task, participants had unlimited response time, and the stimulus changed only after the participant touched the screen.

#### Kanizsa Drag-and-Drop Game

A drag-and-drop matching task (Figure 1d and 1e; Supplementary Tables S3a-S3c) assessed goal-directed use of global illusory contours across three conditions. In the *central faint-colored KIC* and *centered KIC* conditions, participants selected one of four peripheral solid shapes and dragged it to a central target, which was either a faintly colored solid shape or a Kanizsa-induced contour (KIC), respectively. In the *centered solid-shape* condition, a solid shape was presented centrally and participants dragged it from the center to the matching peripheral KIC. The stimulus set comprised nine geometric shapes (including concave and convex forms) designed to discourage local corner-based matching and to emphasize global shape correspondence. Because the solid shapes only approximated the inducer-defined contours, successful matching reflects use of the overall shape rather than precise local alignment. Each solid shape had a corresponding Kanizsa configuration, yielding 18 distinct stimuli presented in five colors (red, green, blue, yellow, purple). On each trial, four peripheral stimuli of identical color but different shapes were displayed in the screen corners, alongside a single central stimulus that was either color-matched or color-mismatched to the periphery.

**Figure 1.**
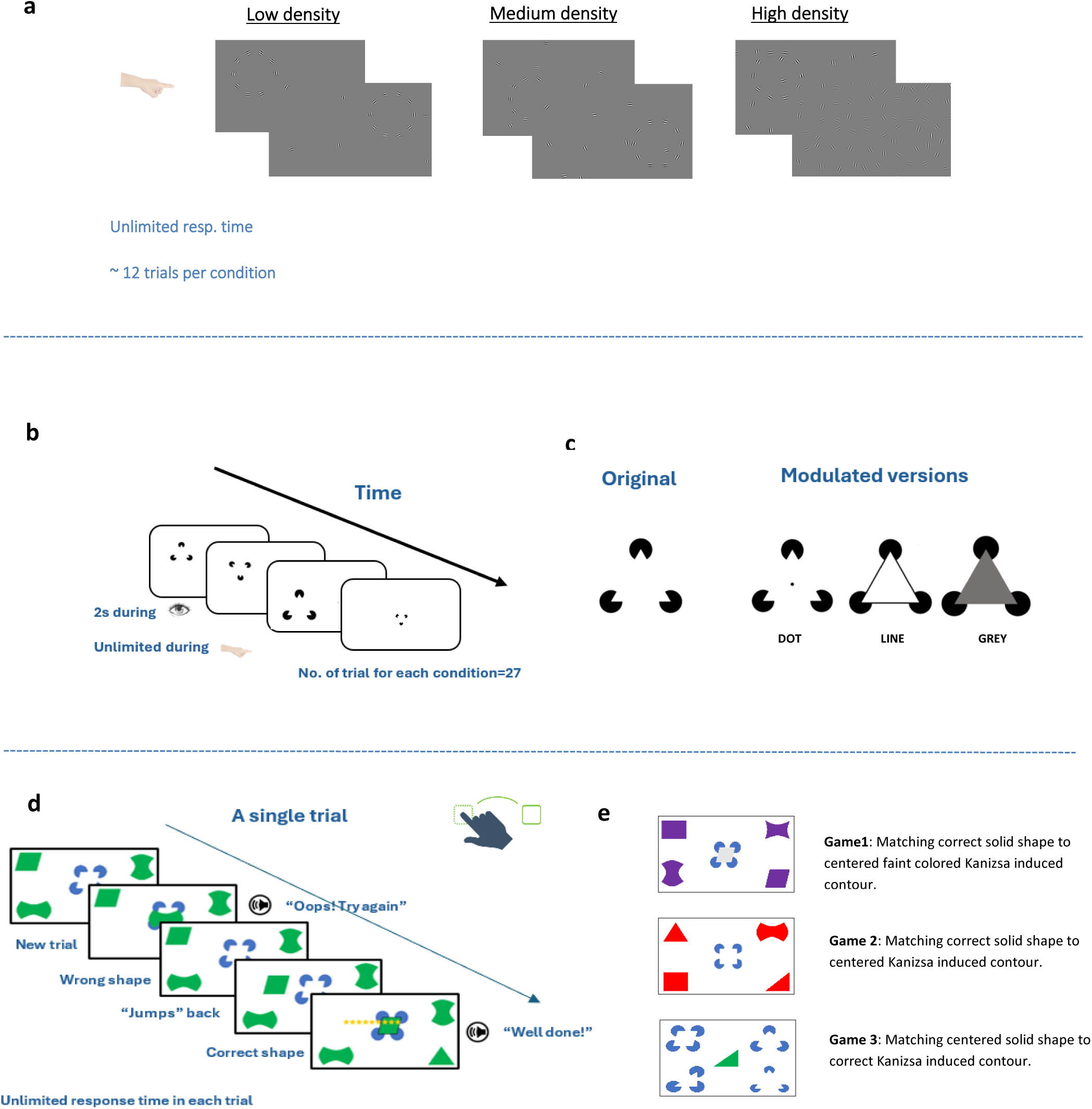
Overview of experimental paradigms and visual stimuli used to assess global contour perception and perception–action dissociations. **(a)** Circular contour (CC) task—experimental paradigm and sample stimuli. Participants were instructed to “point to the circle” under three distractor Gabor density conditions (low, medium, high). The circular target appeared randomly in one of six possible locations. **(b)** Triangular Kanizsa induced contour (KIC) task—experimental paradigm and sample stimuli. Participants were instructed to “point to” or “look at” what appeared on the screen. The KIC appeared randomly in one of four screen quadrants or at the center, in one of three sizes. **(c)** KIC stimulus variants—the original configuration and three modulated versions (Dot, Line, and Grey-luminance). **(d)** Kanizsa touchscreen drag-and-drop game—experimental paradigm and sample stimuli. Participants were instructed to “drag and match the solid shape” to the corresponding global KIC shape. **(e)** Game conditions—three task variants (Game 1, Game 2, and Game 3). See Supplementary Materials for additional details on experimental stimuli. Note: All stimuli shown are adjusted for illustration purposes and do not reflect their exact size, spacing, or on-screen resolution during testing.

### Procedure

Testing occurred in quiet rooms at schools or homes. The first author (HS) conducted all sessions. Participants sat to the left of the experimenter and could move freely; sessions lasted 10–15 minutes and were scheduled once weekly over 3–8 months. Each experimental task began with 8–10 practice trials including demonstration or hand-over-hand guidance if needed and if participant consented. For the CC task, participants were instructed to “point to the circle.” Each condition included approximately 12 trials per density level. For the KIC tasks, the instruction was “point to/look at what appears on the screen”, without explicit instruction regarding the illusory contour with original and modulated conditions presented in separate blocks of approximately 20–24 trials. All participants were first tested on the original triangular KIC in both pointing and eye-gaze, followed by modulated conditions in subsequent weekly sessions. For the Drag-and-Drop game, the instruction was “drag the shape to where it fits,” with each version comprising of 8–12 trials. Rewards (toys, videos, and edible reinforcers) were provided contingent on cooperation and correct responses.

## Data Acquisition and Analysis

### Pointing

#### Circular Contour (CC)

Correct responses were defined as touches landing within 160 px of the target center. Accuracy was computed separately for each level of Gabor distractor density. Within correct responses, touches were further categorized as center pointing (<80 px from the CC center) or frame pointing (≥80 px and ≤160 px from the CC center). Correct pointing was additionally classified as “central Gabor pointing” when the touch occurred within 50 px of a single Gabor element located at the center of the CC (i.e., <100 px from the CC center, to exclude touches on the frame). For all trials, the angular direction of each touch relative to the stimulus center was recorded and circularly normalized within participants. Directional concentration (R) was computed to quantify within-participant angular consistency across trials. For each participant, pointing angles 𝜃 were expressed relative to that participant’s mean direction (𝜃 = 0^∘^ indicates the participant’s preferred direction). Consistency was defined as the mean resultant length from circular statistics:

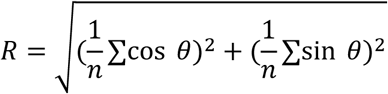

Where n is the number of trials. R=1 indicates perfect alignment of all pointing responses, whereas R=0 indicates random or uniformly distributed pointing directions.

#### Single Kanizsa-Induced Contour (KIC)

For each trial, we computed the Euclidean distance between the participant’s touch location and the centroid of the KIC stimulus. Distances were then normalized such that 0 corresponded to the illusory triangle center and 1 corresponded to the center of the nearest “Pac-Man” inducer. Touches with normalized distances greater than 2.5 were excluded (9.3% on average) because such values reflected aimless or non-task-directed pointing far outside the stimulus region and therefore did not provide interpretable perceptual information. Smaller values indicate more central (global) pointing, whereas larger values indicate pointing near the local inducers. For both the CC and KIC tasks, touches occurring within 200 ms of a previous touch (∼0.6% of all trials, 12 participants, ∼3 trials each on average) were removed to avoid rapid double-taps, which were common in mvASD participants.

### Eye-gaze

#### Single Kanizsa-Induced Contour (KIC)

eye-gaze off-center position was calculated as the Euclidean distance to the center of the KIC stimulus, similar to pointing. Eye-gaze data were analyzed within the 300–700 ms window following stimulus onset.

### Drag-and-Drop motor response

#### Kanizsa Drag and Drop Matching Game

Participants could attempt up to two placements per trial. Scoring rules: (1) Valid placement attempt: an attempt was registered correct/incorrect when the dragged item was released near KIC center (∼0.5cm or 30 pixels center-center distance). If released further away, the solid shape snapped back to its origin and the attempt was not recorded. Across all tested participants, no instances were registered in which shapes were deliberately positioned near the target for comparison prior to a placement attempt. (2) Trial outcome: correct on first or second attempt = Correct; otherwise, Incorrect.

### RCPM Scoring

RCPM raw scores were converted to age-normed percentiles (Raven et al., 1998), interpolated linearly, and scaled to approximate IQ scores (IQ = 100 + z×15), like in Courchesne et al. (2015).

### Subgrouping

K-means clustering (k = 2) reported in Sykes-Haas & Bonneh, 2025 identified two distinct performance subgroups, characterized as ‘high’ and ‘low’ performers. Participants were labeled using pseudonyms beginning with H for high performers (e.g., “Hugo”) and L for low performers (e.g., “Lia”). The same performance classification was applied in the current study to examine subgroup differences in task performance. To explore variability within the Kanizsa-Induced Contour (KIC) tasks, we additionally inspected scatterplots of responses under original and modulated conditions. Clustering was performed using k-means clustering (MATLAB, kmeans), which partitions observations into k clusters by minimizing the within-cluster sum of squared Euclidean distances to cluster centroids. Cluster assignment was based on Euclidean distance in the feature space. The algorithm was run with multiple random initializations, and the solution with the lowest total within-cluster variance was retained. Separate K-means clustering analyses (k = 4 for pointing and k = 3 for eye-gaze data) were conducted to identify participant subgroups showing distinct response patterns to the different stimulus conditions.

### Statistical Analyses

All statistical analyses were conducted using MATLAB R2024a. In the CC task we compared performance within and between groups using independent-samples *t*-tests between typically developing (TD) and minimally verbal autistic (mvASD) participants for (a) the proportion of central Gabor pointing in the Circular Contour (CC) and (b) spontaneous correct center versus frame pointing distributions.

A linear mixed-effects (LME) model was used to evaluate correct responding as a function of Gabor distractor density in the CC task, with density level entered as a fixed effect and participant as a random effect. To assess directional consistency (R) in frame pointing between groups, a non-parametric permutation test was applied to compare the 95 % bootstrap confidence intervals of mean angular directions

During spontaneous responding to KIC normalized off-center pointing and eye-gaze distances performance was analyzed to assess (i) overall differences between groups, (ii) differences across experimental KIC modulation conditions, and (iii) potential interactions between group and KIC modulation condition. This was achieved using a two-way mixed-design ANOVA with repeated measures on the Condition factor. Significant effects were followed up with paired-samples t-tests, which served as post-hoc comparisons between the original and modified KIC conditions (Central Dot, Line, and Grey-luminance).

### Correlational analyses

To control the number of comparisons, we used a hierarchical approach: we first tested associations between descriptive measures (RCPMp, L-VIS, SCQ) and CC indices; only if these were significant would we examine corresponding associations with KIC indices. We then focused on the a priori cross-task question of whether CC global/local spontaneous pointing predicted KIC normalized off-center pointing. For each comparison, effect sizes were computed using Cohen’s *d*, Statistical significance was evaluated at *p* < 0.05 (two-tailed).

## Results

### Circular Colinear Contours (CC)

To examine the ability to detect global colinear contours (CC) under varying levels of local visual noise, Gabor distractor density, participants were asked to point to the circular Gabor-defined contour presented among the distractor elements (Figure 1a). Individual participant’s performance across low-, medium-, and high-density conditions is shown in Figure 2a, and group-level accuracy trends are shown in Figure 2b. Performance accuracy in mvASD participants decreased as distractor density increased, as indicated by a significant negative slope in the LME model p_lme_<0.00005. This result suggests that while mvASD children can detect global contours, their behavioral (i.e. pointing) performance is sensitive to increased local interference.

**Figure 2.**
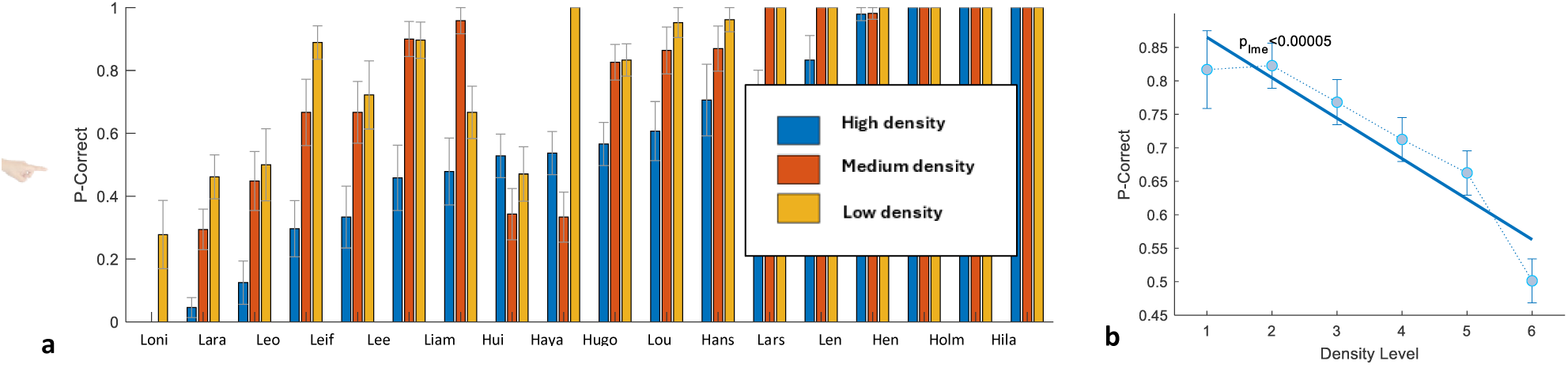
Performance on the contour detection task as a function of Gabor distractor density via pointing among mvASD participants: **(a)** Individual accuracy data (p-correct) are plotted with a bar for each mvASD participant across three distractor density conditions (low, medium, and high) sorted from low to high performance in ‘High density’ condition. Error bars represent ±1 standard error of the mean (SEM). **(b)** Group-level performance as a function of Distractor Density Level, analyzed using a linear mixed-effects (LME) model. The plot shows mean accuracy across participants (±SEM) and the fitted LME regression line, which revealed a significant negative relationship between distractor density and performance accuracy (pₗₘₑ < 0.00005).

To characterize how participants behaviorally detected the CC, we examined their pointing responses characteristics and categorized them as either “center pointing” or “frame pointing” (see Methods and Figure 3). Figure 3a displays mean % trials each experimental group (TDs and mvASD) detected circle either via center pointing or via or frame pointing. Independent sample t-tests revealed that, while both mvASD and TD children successfully detected the CC, mvASD participants pointed significantly more often along the frame of the contour compared with TD peers (Figure 3b, t(37) = -3.62, p <0.001 and Cohen’s d = 1.16, mvASD: ∼30% vs. TD: <10%) and to local Gabor patches within the circle (Figure 3c, t(37)=-2.54, p=0.016 and Cohen’s d=0.81 mvASD: ∼45% vs TD:∼35%), when pointing centrally. TD children showed a higher proportion of center-pointing responses (t(37)=3.84, p<0.001 and Cohen’s d=1.23 TD:>90%). This behavioral pattern indicates that mvASD children may rely more heavily on local salient elements or anchors even when detecting the global structure.

**Figure 3.**
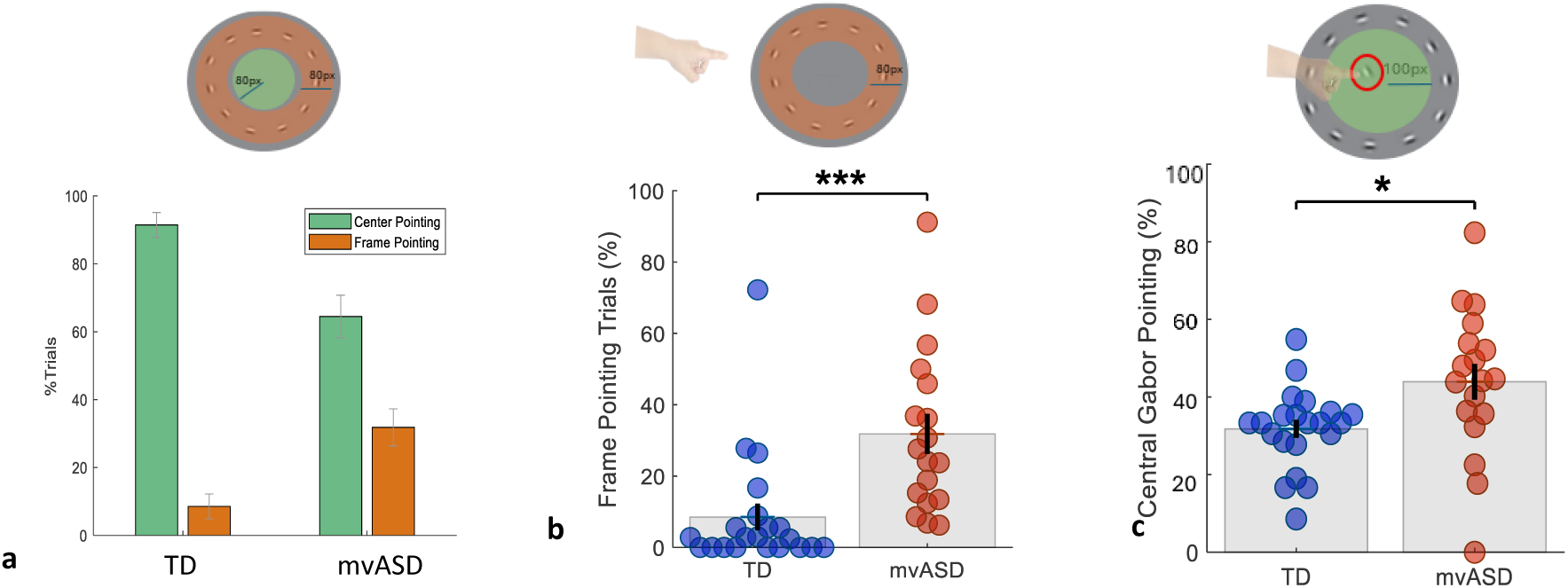
Pointing behavior on Circular contour (CC) in TD and mvASD groups during correct trials: **(a)** Percentage of trials in which participants pointed to the center (green) or the frame of the CC (orange). Error bars represent SEM. Beeswarm plots showing the proportion of correct trials in which each participant **(b)** pointed to the frame of the CC or **(c)** a Gabor patch within the center of the contour CC. Each dot represents one participant; bars indicate group means. Note, whilst successfully detecting the circular contour mvASDs point significantly more often to (local) Gabor patch and on the frame than TDs.

Unexpectedly, mvASD participants displayed spontaneous *localized angular* pointing on the CC frame, typically confined within half of the contour or less, irrespective of CC location on the screen (i.e. 6 randomized locations, Figure 1a and Figure 4). As previously noted, TD participants’ spontaneous pointing was directed toward the CC center. To determine whether the localized angular pointing among mvASD reflected a general *behavioral* strategy rather than perceptual differences between groups, TD participants were additionally tested under an instructed condition in which they were asked to point anywhere on the CC frame. Normalized polar heatmaps of spontaneous angular frame pointing distributions for mvASD and TD children are shown in Figure 4. Each polar plot represents the distribution of normalized pointing angles across all trials, where 0° corresponds to each participant’s own mean direction. Both groups’ responses were largely restricted to a half-circle sector, indicating consistent angular localization relative to each participant’s preferred pointing location. Directional concentration (Figure 4), quantified by the mean resultant length (*R*), was moderately higher in TD children (*R* = 0.62 and 95% CI: 0.53 – 0.72) than in mvASD participants (*R* = 0.55, 95% CI: 0.45 – 0.65). The observed difference in mean *R* (ΔR = –0.070, CI 95%: -0.206 to 0.064) did not reach statistical significance (permutation test, *p* = 0.34). Thus, although TD children showed slightly greater within-participant consistency in their pointing direction, both groups exhibited comparable degrees of localized angular repetition.

**Figure 4.**
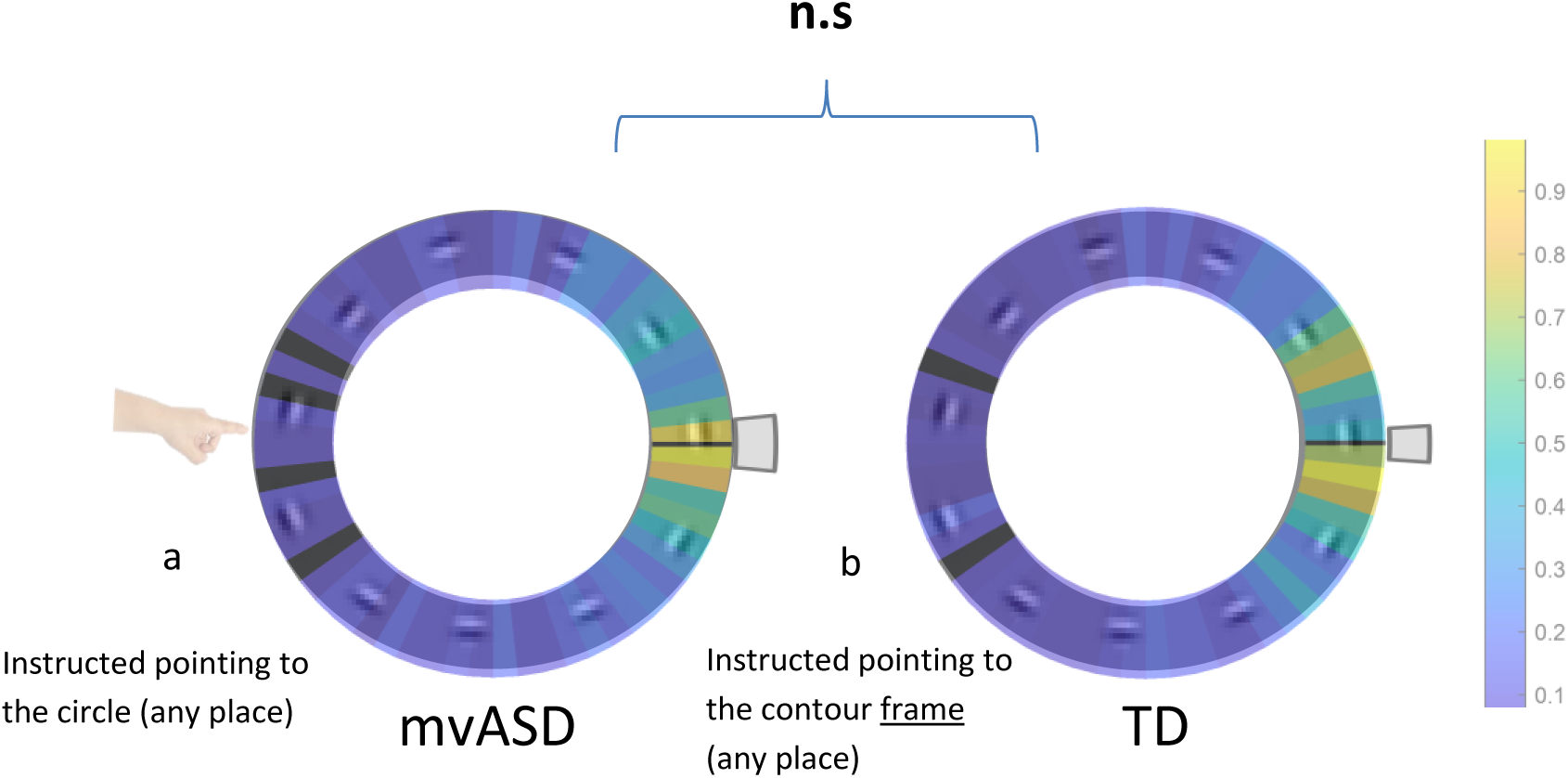
Normalized polar heatmaps of spontaneous angular pointing distributions on the circular contour (CC) frame for mvASD and TD children. Each polar plot shows the distribution of normalized spontaneous pointing angles across all trials, **(a)** mvASD children’s data reflect spontaneous frame-pointing, whereas **(b)** TD data reflect instructed frame-pointing, where 0° corresponds to each participant’s individual mean direction. Color intensity on the ring represents the relative frequency of pointing responses (yellow = higher density, blue = lower density). The black radial line marks the group mean direction, and the gray arc surrounding it denotes the 95 % bootstrap confidence interval for that mean, estimated from resampled trials. A narrower confidence interval indicates greater certainty and directional consistency across responses. Note, both groups show responses confined largely within a half-circle, but the TD group demonstrates a slightly higher concentration (R = 0.62 vs 0.55) and narrower confidence interval, but this difference in non-significant (via a permutation test).

### Kanizsa Induced Contours (Pointing)

In the Kanizsa Induced Contour (KIC) task (Figure 1b and 1c), participants were instructed to “point to what you see on the screen.” A mixed-design ANOVA with group (mvASD vs. TD) as a between-subjects factor and modulation condition (no modulation vs. three modulation types) as a within-subjects factor revealed a significant main effect of group, *F*(1,36)=12.27, *p*=0.001, ηp²=0.25, no main effect of modulation, *F*(3,108)=1.45, *p*=0.23, ηp²=0.04, and a significant group × modulation interaction, *F*(3,108)=8.51, *p*<0.001, ηp²=0.19, indicating that modulation effects differed between the mvASD and TD groups.

Post-hoc comparisons using independent-samples *t*-tests showed that, in the original (unmodulated) KIC condition, mvASD participants pointed significantly closer to the local ‘Pac-Man’ inducers, whereas TD participants more often pointed toward the illusory center (*t*(42)=−5.30, *p*<0.0001, Cohen’s *d*=1.60; Figure 5a,b).

Within the mvASD group, post-hoc paired-samples *t*-tests revealed that KIC cue modulation (collapsed across Grey -luminance, Central Dot, and Line conditions; Figure 5d) led to a significant shift from off-center pointing toward the illusory center (*t*(20)=2.33, *p*=0.037, Cohen’s *d*=0.53). Analyses of individual modulation conditions (Figure 5c) showed significant center ward shifts during luminance modulation (*t*(20)=3.02, *p*=0.030, Cohen’s *d*=0.66), Central Dot modulation (*t*(20)=2.06, *p*=0.050, Cohen’s *d*=0.45), and line modulation (*t*(20)=2.25, *p*=0.046, Cohen’s *d*=0.53).

K-means clustering (*K*=4) further revealed marked individual variability, with a subset of mvASD participants showing increased global contour selection under enhanced cueing in the modulated KIC conditions (Figure 5e). Overall, these results indicate reduced spontaneous behavioral use of global illusory contours in mvASD participants, alongside improved utilization of global contour information when local cues are explicitly enhanced (Figure 5a–e).

**Figure 5.**
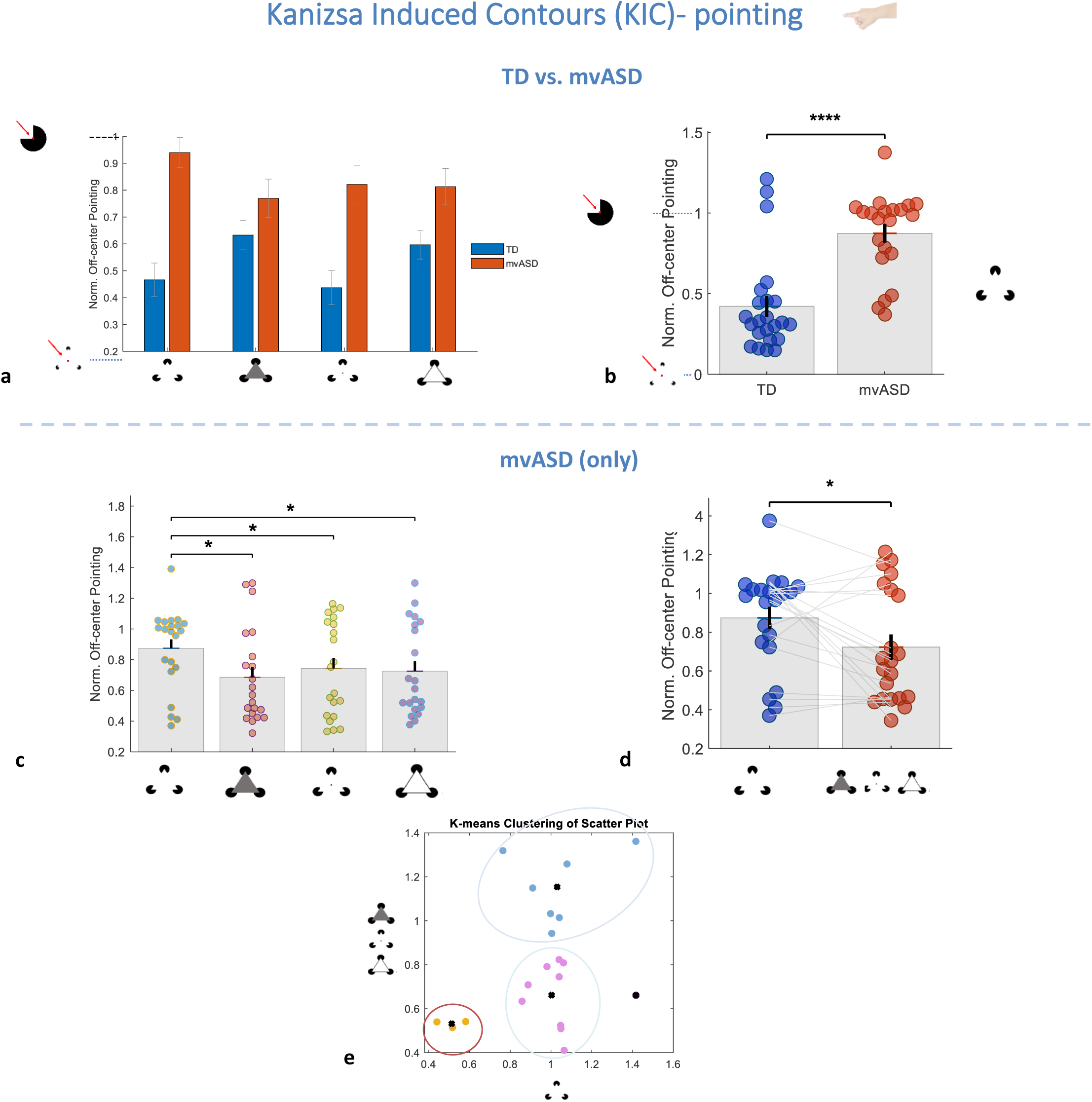
Normalized spontaneous off-center pointing on Kanizsa induced contours (KIC) in original and modulated (grey-luminance, central dot and line) conditions among children with mvASD and TD: **(a)** Bar graph showing group means across all conditions. **(b)** Beeswarm plot showing individual data points and group means for TD and mvASD in the Original triangle KIC condition (p<0.0001), **(c)** – **(e)** mvASD data only: **(c)** Beeswarm plots comparing Original KIC condition with each of the modulated KIC conditions seperately, **(d)** comparison of the original KIC condition with all modulated KIC conditions combined and **(e)** Diagonal scatterplot where each circle represents one mvASD participant’s spontaneous off-center pointing in original and modulated KIC conditions. Colored circles indicate clusters identified using K-means clustering (K = 4). Note, predominantly local Pac-Man inducer pointing among mvASD in the absence of contour-enhancing modulations, indicating reduced spontaneous behavioral use of global contour.

### Kanizsa Induced Contours (Eye-Gaze)

Eye-gaze data (Figure 6a-f) revealed a pattern paralleling pointing behavior. A mixed-design ANOVA with group (mvASD vs. TD) as a between-subjects factor and modulation condition as a within-subjects factor showed a significant main effect of group, *F*(1,38)=18.97, *p*<0.001, ηp²=0.33, and a significant main effect of modulation, *F*(3,114)=3.71, *p*=0.014, ηp²=0.09, with no group × modulation interaction, *F*(3,114)=0.24, *p*=0.87, indicating that modulation influenced gaze behavior similarly in both groups. Specifically, mvASD participants’ fixations were primarily directed toward local KIC inducer regions, whereas TD children more often fixated nearer the illusory center (*t*(40)=−4.41, *p*<0.0001, Cohen’s *d*=1.37; Figure 6a,b).

Within the mvASD group, a post-hoc paired-samples *t*-test comparing the original KIC condition with the combined modulation conditions (luminance, central dot, and line) revealed a significant shift in eye gaze toward the illusory center (*t*(18)=2.23, *p*=0.038, Cohen’s *d*=0.62). Paired comparisons between the original condition and each individual modulation condition did not reach significance. Histogram and scatter analyses further showed that mvASD gaze distributions shifted toward perceptually global centers when modulating cues were introduced. K-means clustering (*K*=3) identified subgroups differing in global versus local fixation strategies. Together, these findings suggest that mvASD participants spontaneously prioritize salient local regions over inferred global contours, consistent with reduced weighting of top-down global guidance.

**Figure 6.**
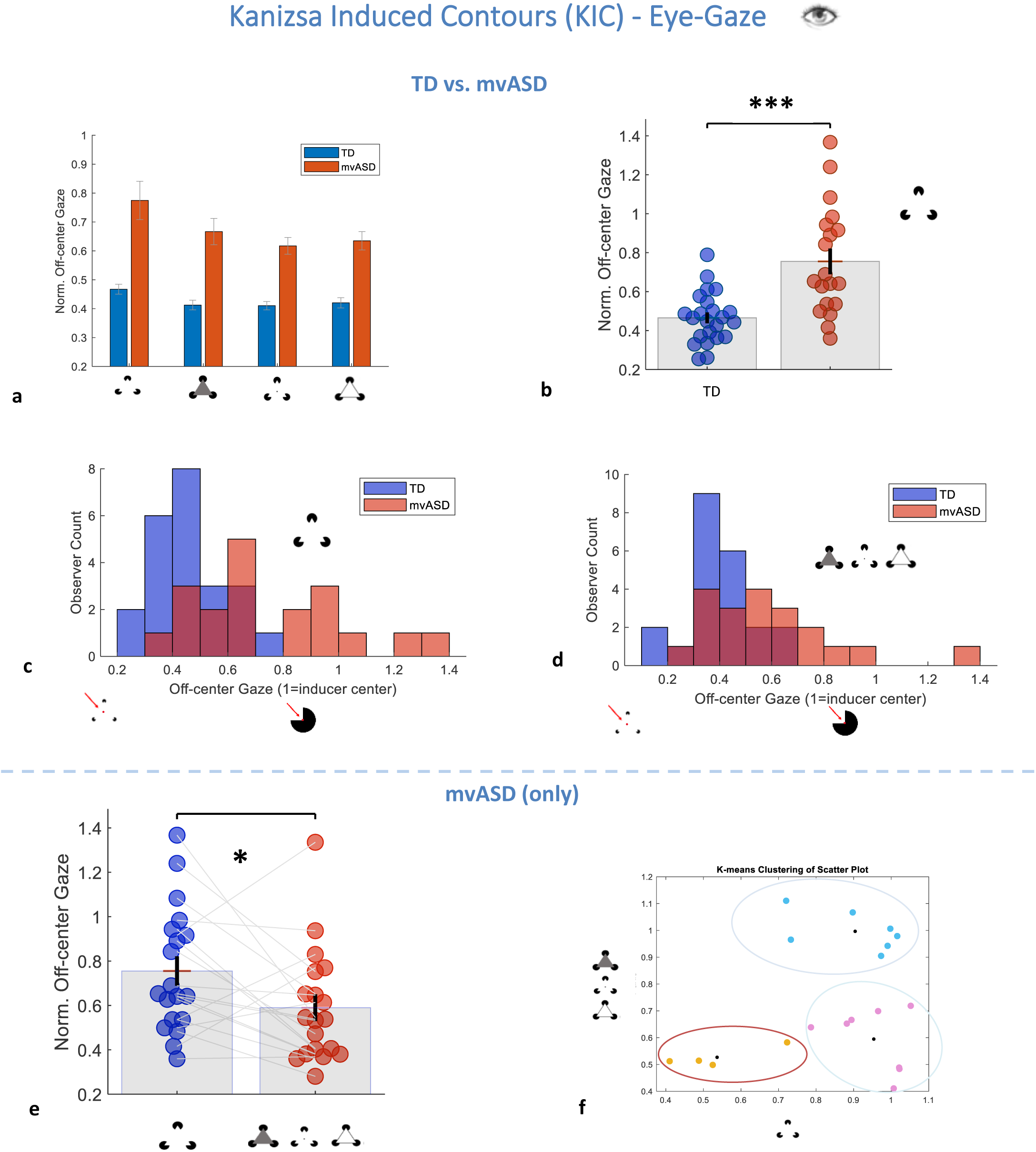
Normalized spontaneous off-center eye-gaze on triangular Kanizsa induced contours (KIC) in original and modulated (grey-luminance, central dot and line) conditions among children with mvASD and TD: **(a)** Bar graph showing group means across all KIC conditions. **(b)** Beeswarm plot showing individual data points and group means for TD and mvASD in the Original KIC condition (p<0.0001), **(c–d)** Histograms showing observer counts as a function of off-center gaze position (0 = illusory center, 1 = inducer center) for both groups in original and modulated KIC version, respectively. **(e)** and **(f)** mvASD data only: **(f)** comparison of the original KIC condition with all modulated KIC conditions combined and **(d)** Diagonal scatterplot where each circle represents one mvASD participant’s spontaneous off-center eye-gaze in original and modulated KIC conditions. Colored circles indicate clusters identified using K-means clustering (K = 3). Note, similar to the pointing data, mvASD participants showed primarily local, near-inducer gaze when contour cues were unmodulated, suggesting reduced spontaneous selection of the global contour during eye-gaze.

### Kanizsa Drag-and-Drop Game

To evaluate whether mvASD participants can explicitly match a variety KIC shapes to corresponding solid shapes, a drag-and-drop game was administered (Figure 1d and 1e). 90% of participants performed above chance in at least one of the three game versions, with best performance observed in Game 3 (Figure 7a, solid central shaped dragged to correct 1 of 4 KIC options). Most ‘high’ mvASD performers (pseudonyms beginning with ‘H’; Sykes-Haas & Bonneh, 2025) achieved near-ceiling accuracy across all game versions, indicating preserved perception of illusory shapes despite altered spontaneous pointing and gaze behavior in KIC and CC tasks. These results demonstrate that mvASD participants can perceive and utilize global contours when task demands emphasize goal-directed matching rather than spontaneous orienting.

**Figure 7.**
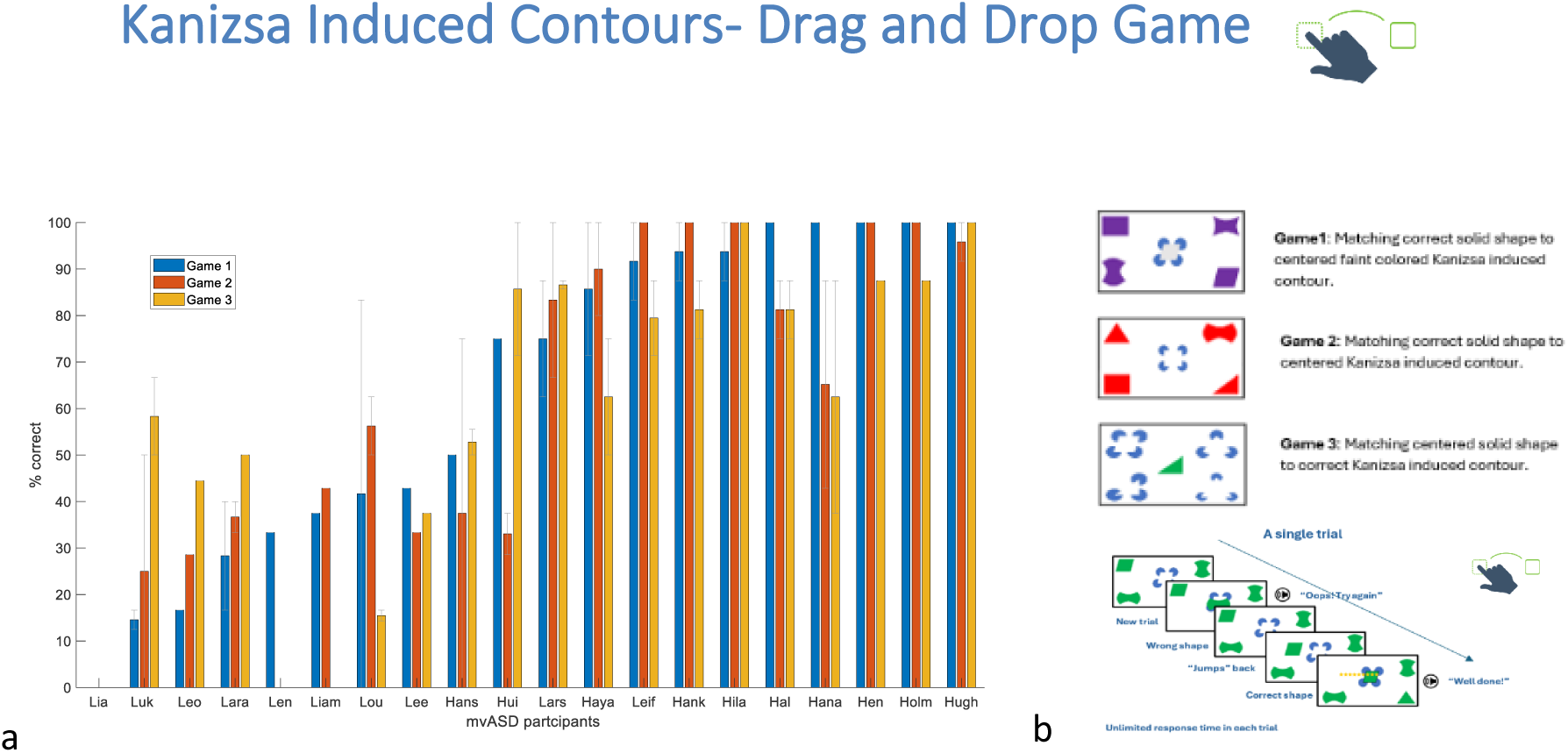
Performance on the Kanizsa Induced Contour (KIC) drag-and-drop game among mvASD participants. **(a)** Bar graph showing individual mean percent-correct performance across the three KIC game conditions. **(b)** Schematic illustration of the three KIC game conditions and trial flow. Note, all mvASD participants performed above chance in at least one KIC game condition, primarily in Game 3. High-performing participants (pseudonyms beginning with ‘H’; Sykes-Haas & Bonneh, 2025) performed above chance across all KIC game conditions, majority reached or approached ceiling performance. Finding suggest successful perception of global illusion and behavioral <u>use</u> of global contour.

### Correlations

#### CC pointing and KIC off-center pointing (Figure 8a-c)

In the mvASD group normalized off-center pointing distance in KIC was significantly associated with CC pointing indices: frame pointing (r = 0.69, p = .0033; Figure 8a), centralized Gabor pointing (r = 0.55, p = .027; Figure 8b), and Gabor-or-frame pointing (r = 0.55, p = .028; Figure 8c). Thus, higher CC local/element-based pointing (frame and central-Gabor indices) was associated with more off-center KIC pointing, that is, pointing closer to Pac-Man inducer centers rather than the illusory contour center.

#### Eye-gaze and pointing cross-size consistency (Figure 8d-g)

Normalized off-center <u>eye-gaze</u> distance was strongly correlated between KIC Size 1 and KIC Size 3 in both groups (TD: r = 0.83, p < .00005; Figure 8d; mvASD: r = 0.92, p < .00005; Figure 8e). Normalized off-center <u>pointing</u> distance also showed strong cross-size consistency (TD: r = 0.97, p < .00005; Figure 8f; mvASD: r = 0.82, p < .00005; Figure 8g). Together, these results indicate strong cross-size consistency in spontaneous eye-gaze and pointing behavior during KIC.

#### Descriptive measures and CC pointing

Descriptive measures (RCPMp, L-VIS, SCQ) were not associated with CC pointing indices (all ps > .05); therefore, we did not test additional descriptive-measure correlations with KIC indices.

**Figure 8.**
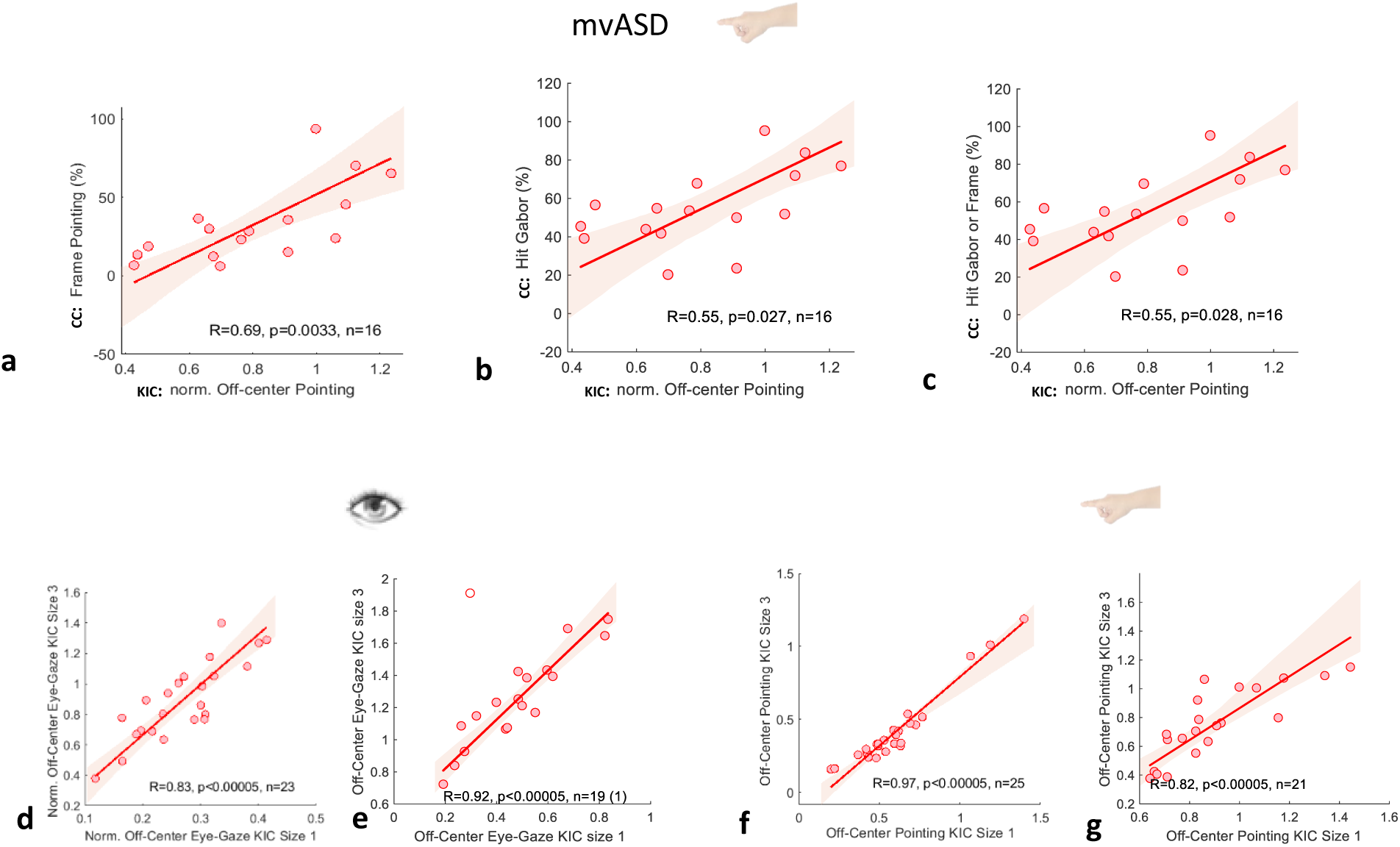
Cross-task and cross-size correlations of pointing and eye-gaze indices in mvASD and TD groups. Cross-task correlations between spontaneous pointing to Kanizsa-induced contours (KIC) and pointing strategies in the circular contour (CC) task. The x-axis shows normalized off-center pointing distance in the KIC task; the y-axes show CC pointing indices: **(a)** frame pointing (%), **(b)** centralized Gabor pointing (%), and **(c)** Gabor-or-frame pointing (%). Cross-size correlations between KIC Size 1 and KIC Size 3 are shown for pointing and eye-gaze behavior: **(d)** TD eye-gaze, **(e)** mvASD eye-gaze, **(f)** TD pointing, and **(g)** mvASD pointing. Pearson correlations are reported as r (two-tailed p values); n = 16. Solid lines indicate least-squares linear fits.

## Discussion

In the present study, we examined spontaneous pointing and eye-gaze responses as well as goal directed behavior to global form cues in minimally verbal autistic (mvASD) and typically developing (TD) children. We probed two illusory-contour paradigms: Circular contours (CC) and Kanizsa illusory contours (KIC). We quantified spontaneous pointing in CC during increasing Gabor density conditions. During KIC we quantified spontaneous pointing and eye-gaze during original triangular Kanizsa illusion as well as during modulated versions accentuating the central triangle. We assessed the effect of cue modulations on response localization, and compared spontaneous performance with a goal-directed KIC drag-and-drop task designed to require increased weighting of global form and reduce the utility of local matching.

### Circular Contours (CC) local/global pointing indices in TD and mvASD

During successful detection of CC mvASD children showed a higher proportion of pointing responses directed toward stimulus components (frame and Gabors). Overall they pointed to frame ∼ 30% of trials and central Gabor elements ∼45% of their central pointing trials unlike TDs who consistently spontaneously pointed centrally (Figure 3a-c). In addition, mvASD frame pointing exhibited a global object-centered pattern: After normalizing responses to the CC coordinate frame, responses tended to fall on one half of the circle regardless of the CC’s position on the screen (six alternative locations, Figure 4a). TD children also showed object-centered frame pointing to half the circle, but this pattern was only observed when they were explicitly instructed to point to the CC frame (Figure 4b).

### Kanizsa Illusory Contours (KIC) local/global (pointing) indices in TD and mvASD

In KIC, like in CC, mvASD children showed more localized spontaneous pointing (Figures 5a-e) and eye gaze (Figures 6a-f) near KIC Pac-Man inducers compared with TD children. Within mvASD, the location of spontaneous pointing (Figure 5a, 5c, 5d and 5e) as well as eye-gaze (Figure 6a, 6c, 6d and 6f) shifted towards the center during cue modulations that were applied to accentuate central global triangle in the KIC displays (luminance, central dot and line). During the Kanizsa drag and drop game KIC were constructed with curvature to increase weighting of global percept. Most (∼90%) of mvASD children succeeded above chance in at least one drag and drop KIC task condition and more than half performed at or near ceiling. (Figure 7a).

### Evidence consistent with access to global percepts

Together, these findings suggest that while global contour perception was present in both groups during CC and KIC displays, mvASD children’s spontaneous behavior placed relatively greater weight on local and/or salient elements or anchors than TD’s. Across tasks, several observations were consistent with access to global structure. First, object centered pointing towards one half of the CC, evident in both TDs and mvASD indicate global percept detection and whilst central Gabor pointing and frame pointing are “localized behaviors” participants successfully detected global CC percept, also with increase in Gabor density. Second, during KIC tasks, manipulations that increased the visibility of the central triangle region were accompanied by shifts in both pointing and gaze toward the global structure’s center. Of particular interest is the central dot modulation that maintain the illusory contour whilst simultaneously accentuating its global presence. Finally, mvASD children succeeded in a goal-directed KIC drag-and-drop task in which KIC were constructed with curvature to eliminate reliance on local feature matching. In this setting, performance remained above chance, more than half performing at or near ceiling, indicating that task-relevant information was available for selecting the correct global KIC correspondence.

### Cross-task correlational findings

Across mvASD participants, localized (frame or Gabor) pointing in CC was associated with off-center (localized Pac-Man) pointing in KIC (Figure 8a-c) suggesting a cross-paradigm relation in response localization. We also found strong within-participant cross-size consistency: off-center eye-gaze was strongly correlated across KIC sizes (Figure 8d-e), and off-center pointing was likewise strongly correlated across sizes (figure 8f-g), in both mvASD and TD participants. These findings indicate each localization index is stable across stimulus scale and experimental paradigms and strengthens stable individual response patterns. In contrast, the degree of localized responding was not explained by available descriptive measures (LVIS, RCPMp or SCQ) in this dataset. Although the sample size limits inference from null associations, the robust and stable response patterns in these indices across participants suggest that spontaneous pointing and eye-gaze may reflect systematic differences in underlying perceptual and response-selection processing in mvASD that are not captured by LVIS, RCPMp, or SCQ.

### What could account for atypical local pointing in mvASD?

Although our findings indicate that global processing is present in mvASD, they still present with spontaneous bias towards local elements . These biases could be accounted for by accounts of weaker or potentially slower global inferencing (Nayar et al., 2017; Booth and Happe, 2018, Frith and Happe, 1994, Van der Hallen et al., 2015). It may also be explained as a style and a disinclination to report globally (Koldewyn et al., 2013), as they do succeed in reporting on the global within instructed tasks. In this context, our finding of TDs’ ‘disinclination’ to point ‘locally’ on the frame is notable.

Nevertheless, we advance an alternative account: the same pattern may primarily reflect atypical response selection under ambiguity, when multiple candidate targets compete, rather than perception alone (Robertson and Baron-Cohen, 2017). Across CC as well as KIC, several potential behaviors, i.e., affordances (Gibson, 1979; Cisek, 2007), are concurrently available for selection (center, frame, Gabor, Pac-Man). In TD children, these alternatives appear to be downregulated such that global center-pointing becomes the predominant selected affordance, whereas in mvASD multiple candidate targets may remain concurrently available. Robertson and Baron-Cohen (2017) proposed that differences in canonical micro-circuitry or neural motifs in autism could contribute to diverse phenomena spanning language (homographs; Frith et al., 1983; D’souza et al., 2016), perception, and social cognition (i.e. theory of mind, Baron-Cohen et al., 1985). They suggest that autism involves disruption in ambiguity resolution, normally supported by reciprocal inhibitory competitive interactions between neural populations vying for representational dominance, consistent with evidence from binocular rivalry paradigms by Robertson and colleagues (Spiegel et al., 2019, see also Rubenstein and Merzenich, 2003 for E/I imbalance in autism). This emphasis on competition also aligns with the Affordance Competition Hypothesis (Cisek, 2007), supported by neural population evidence for competition prior to response selection in premotor cortex (Cisek and Kalaska,2005).

In our tasks, spontaneous pointing and eye-gaze do not enforce a single “correct” response target; TD children nevertheless respond relatively consistently, whereas mvASD pointing behavior suggests that multiple response sets/affordances may remain concurrently available. Previous findings, unrelated to the local-global dissociation also show response selection via pointing/touch is challenging for individuals with mvASD and may not reflect true competence (e.g. during word recognition, Ellert et al., 2025 and oddball detection, Sykes-Haas and Bonneh, 2025, WISC-IV assesment, Courchesne et al., 2015). However, we also find that visual saliency and increased deviance facilitate pointing/touch performance on oddball tasks with otherwise identical (cognitive) requirements. Furthermore, when successfully detecting global triangular Kanizsa oddball among square kanizsa distractors, some mvASD choose to point locally. Notably, when alternative response contexts (to pointing) are offered, often more tangible, where the child can touch or move/drag and physically manipulate response materials, performance in mvASD appears more regulated and goal-directed, consistent with preserved competence (e.g. RCPM puzzle vs booklet, drag and drop KIC game, Children’s embedded figures and visual search reported via physical placement; Courchesne et al., 2015 and 2019). We therefore suggest that these more tangible/structured contexts may increase the strength of the task-relevant affordance for selection, reducing ambiguity by increasing salience/deviance, providing richer online feedback, reducing memory or instruction demands, and changing engagement, thereby biasing selection toward the intended target. Thus, the question may not be *what* they perceive, but *how* they use the visual signal for behavior selection.

### Limitation and future directions

These findings should be considered in light of a few constraints. First, the sample size, particularly for cross-task correlations and null associations with descriptive measures, limits the strength of conclusions about individual differences. Second, future work would benefit from richer participant characterization (e.g., standardized adaptive behavior measures), which would help clarify whether stable response-localization patterns relate to broader functional profiles and support needs. Several directions follow naturally from the current design. One is to further develop goal-directed “global completion” tasks that require selecting global form through drag-and-drop or construction-style responses, conceptually paralleling ambiguity-resolution paradigms in language (e.g., homographs), and extending from object-level to scene-level contexts. A second is to use parametric stimulus manipulations to independently vary local salience (e.g., inducer contrast/deviance) and global cue strength (e.g., contour coherence), while distinguishing “central” responses that land on discrete local anchors from those that do not.

More broadly, the present findings motivate a shift in emphasis from asking whether mvASD is “local” or “global” to specifying the conditions under which global information is expressed behaviorally. On this view, global visual information may be available, while spontaneous action selection favors local anchors when multiple response targets compete. This framing yields testable predictions: as ambiguity is reduced and the global affordance is strengthened, via cueing, feedback, or response structure, response localization should shift systematically toward global centers within mvASD individuals.

## Supporting information

Supplemental Table S1 and S2 CC and KIC

Appendix1 Descriptive measures

## Notes

### Competing Interest Statement

The authors have declared no competing interest.

### Funding

- Israel Science Foundation: Grant No. 657/21

