## Supplemental TableS3 for "Minimally verbal children with autism may see the global, but point local: A behavioral and eye-tracking study in visual perceptual processing"

### Supplementary Tables: Drag-and-Drop Stimuli

#### Table S3a. Solid Shape Properties

| Shape ID  (blue color for illustration only) | Bounding Box  (cm, H ×W) | Bounding Box  (px, H ×W) | Feret (px) | Feret (cm) | Feret (deg) |
| --- | --- | --- | --- | --- | --- |
| 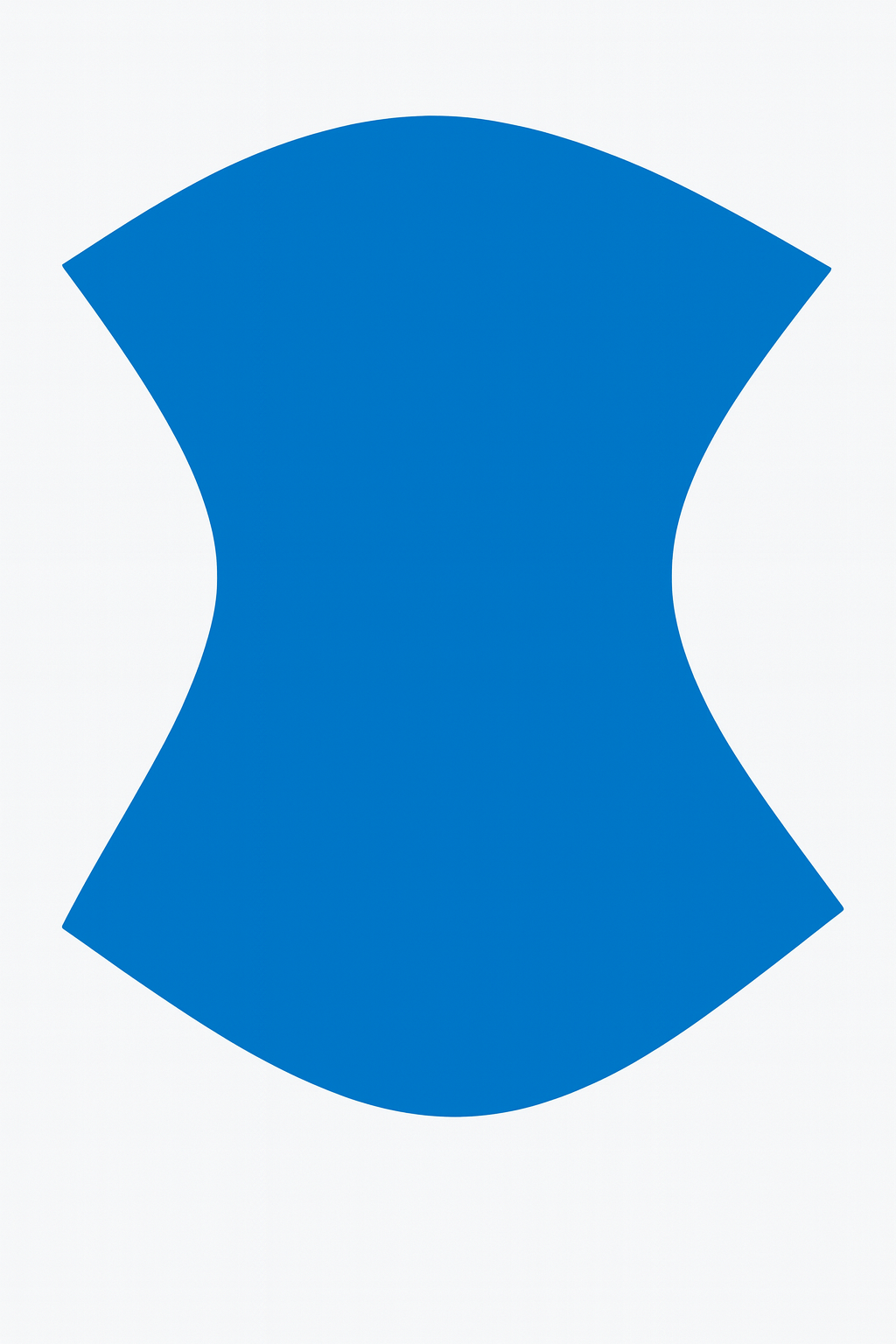1 | 6.0 x 3.5 cm | 371.6 x 217 px | 371.6 px | 6.0 cm | 5.72 ˚ |
| 2 | 4.8 x 4.8 cm | 297 x 297 px | 297 px | 4.8 cm | 4.58 ˚ |
| 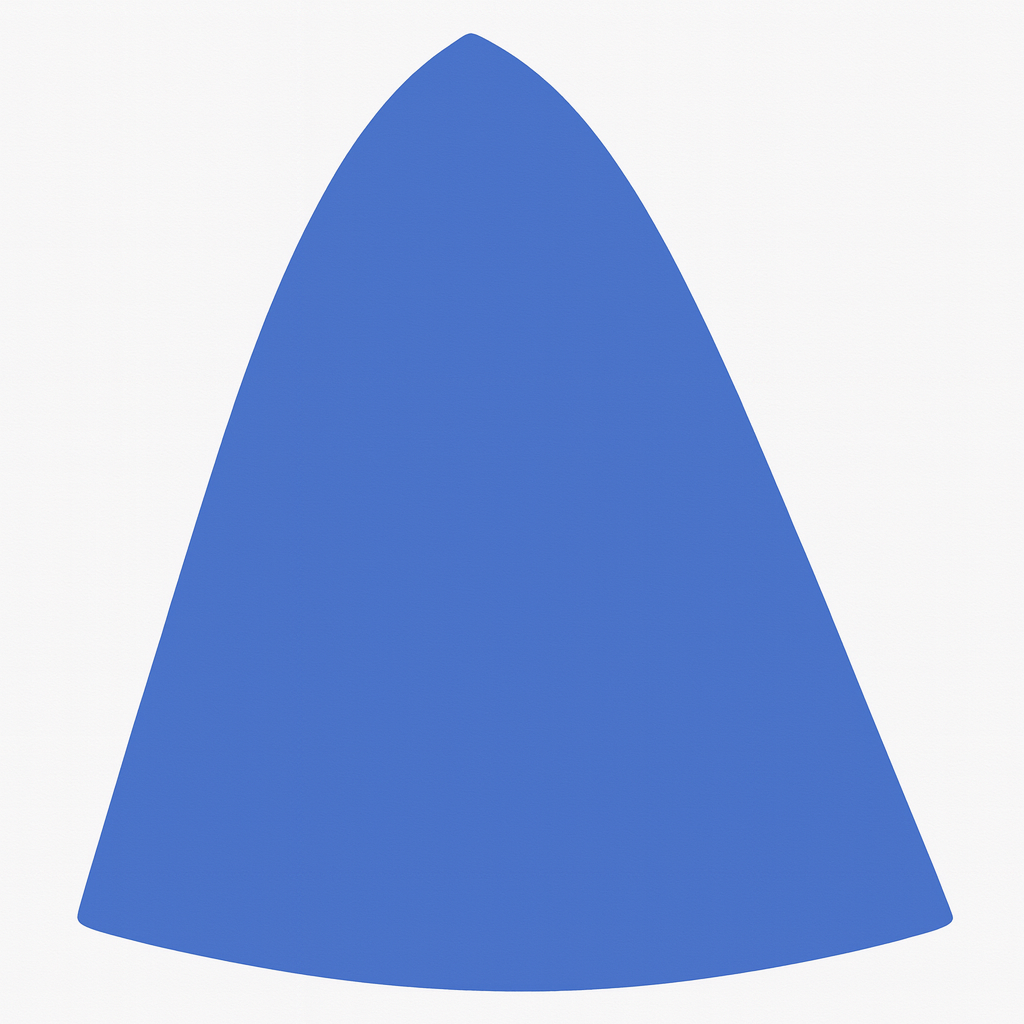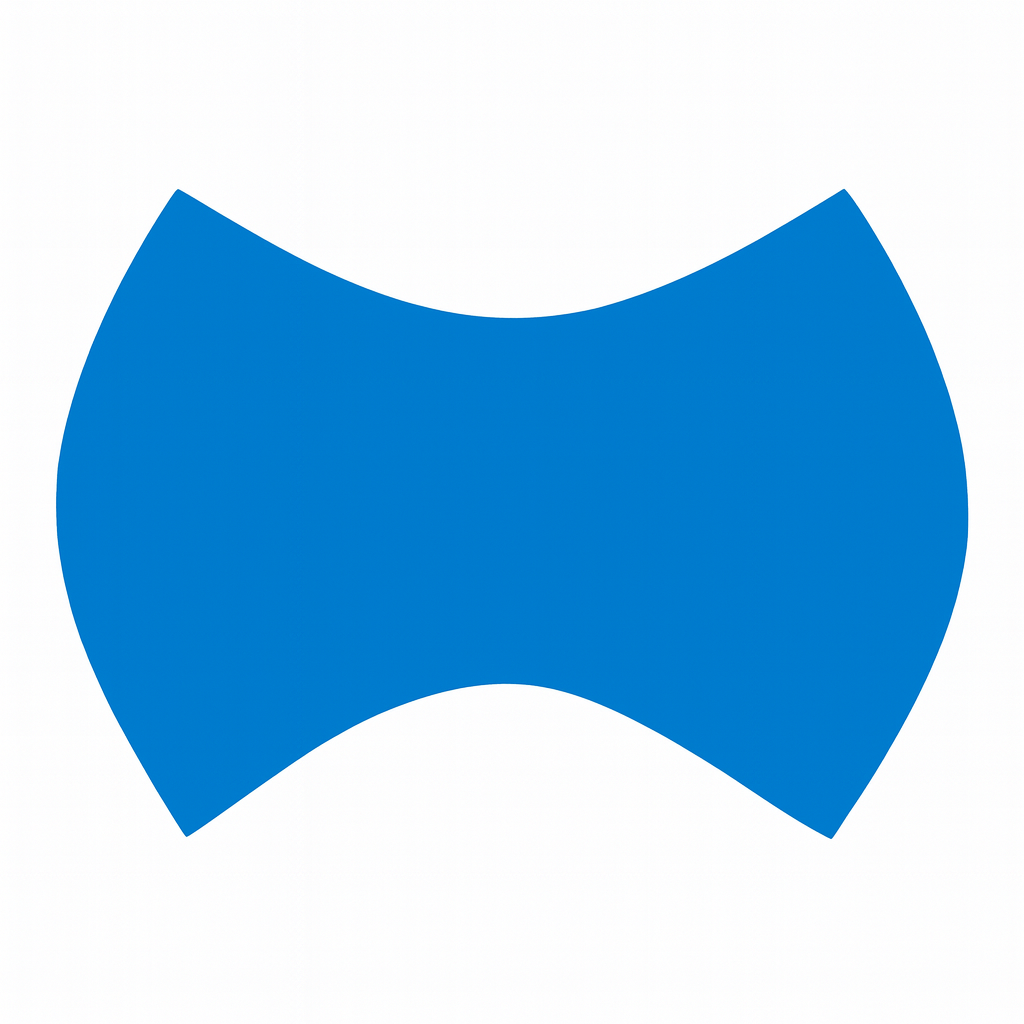3 | 3.5 x 6.0 cm | 217 x 371.6 px | 371.6 px | 6.0 cm | 5.72 ˚ |
| 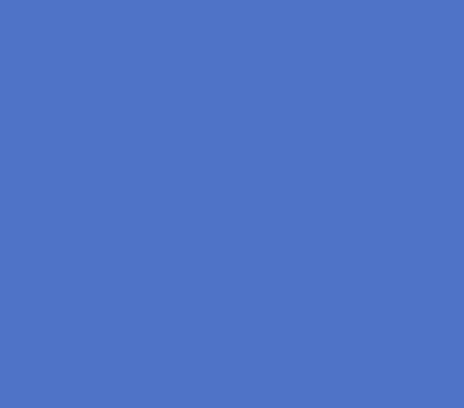4 | 3.5x4.1 cm | 217 x 254.2 px | 371.6 px | 6.0 cm | 5.72 ˚ |
| 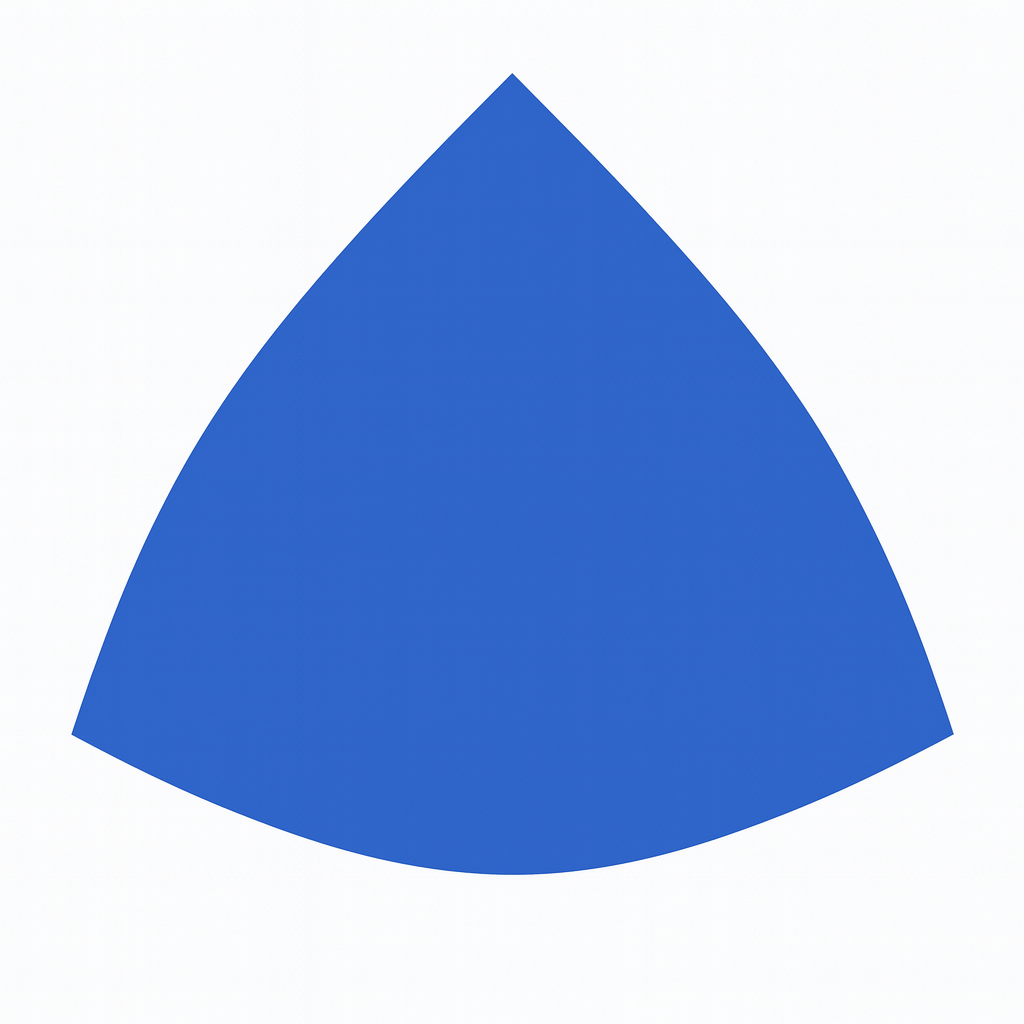5 | 3.5 x 4.5 cm | 217 x 278.7 px | 278.7 px | 4.5 cm | 4.30 ˚ |
| 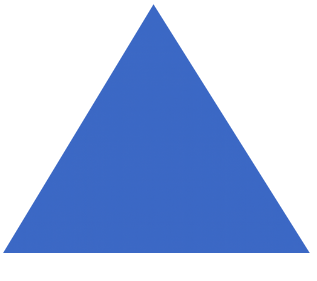6 | 3.5 x 4.7 cm | 217 x 291 px | 291 px | 4.7 cm | 4.49 ˚ |
| 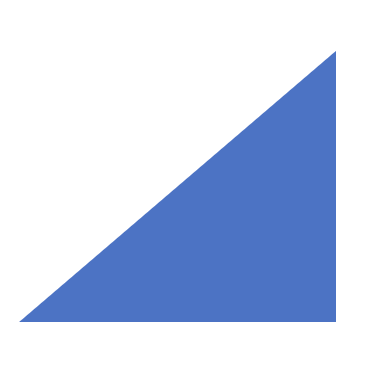7 | 3.5 x 4.5 cm | 217 x 278.7 px | 371.6 px | 6.0 cm | 5.72 ˚ |
| 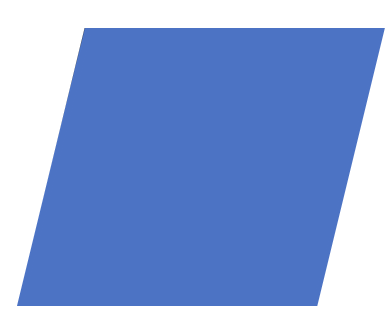8 | 3.5 x 4.7 cm | 217 x 291 px | 371.6 px | 6.0 cm | 5.72 ˚ |
| 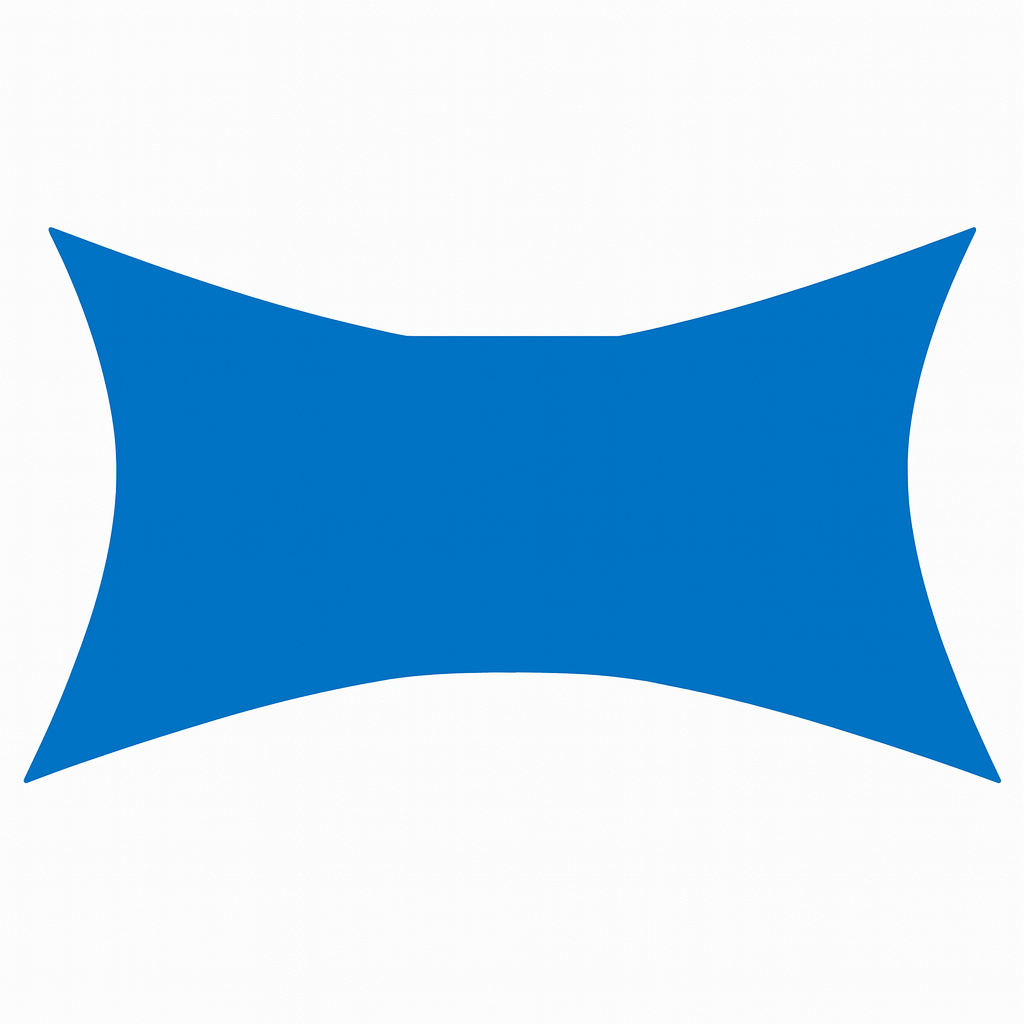9 | 3.8 x 4.7 cm | 217 x 291 px | 371.6 px | 6.0 cm | 5.72 ˚ |

#### Table S3b. Kanizsa (KIC) Inducer Geometry

| KIC ID | Inducers  (n) | Radius (cm / deg) | Opening Angle (deg) of PacMan Inducer | Inter- inducer center spacing (cm / deg) |
| --- | --- | --- | --- | --- |
| 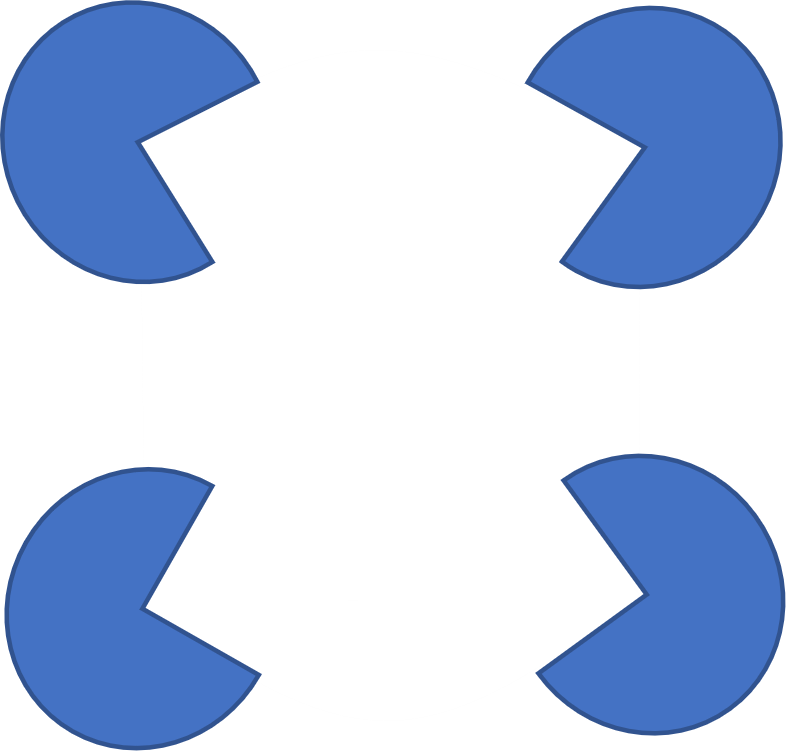1 | 4 | 1.1 cm / 1.1˚ | 90˚ | 4.2 cm (4.01°) ×2  3.3 cm (3.15°) ×2  5.5 cm (5.24°) (Feret) |
| 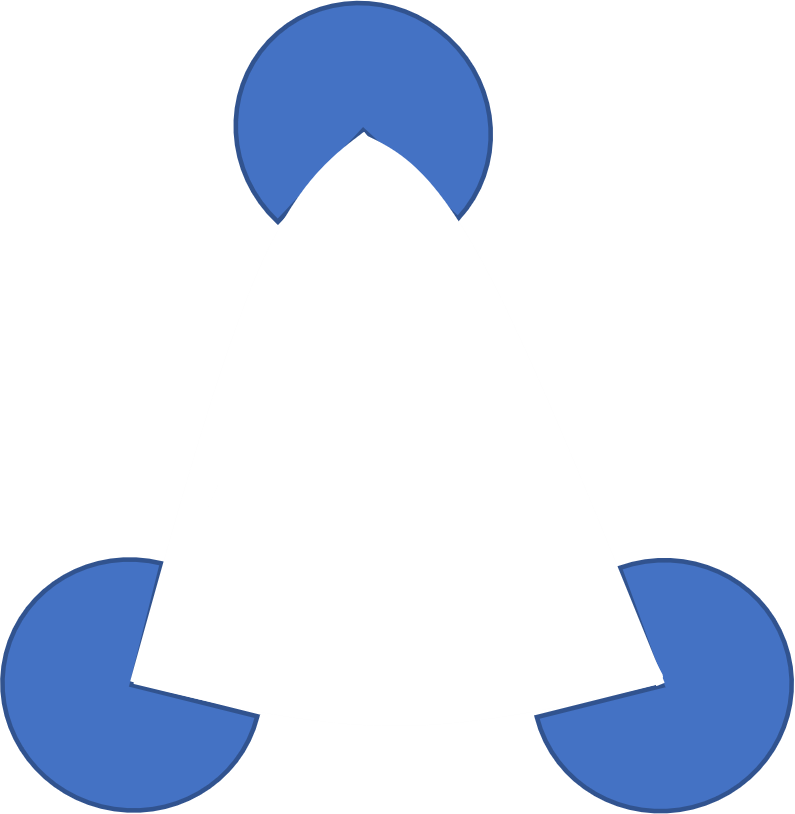2 | 3 | 1.0 cm / 1.0˚ | 90˚ | 4.5 cm (4.29°) ×1  5.0 cm (4.77°) ×2 |
| 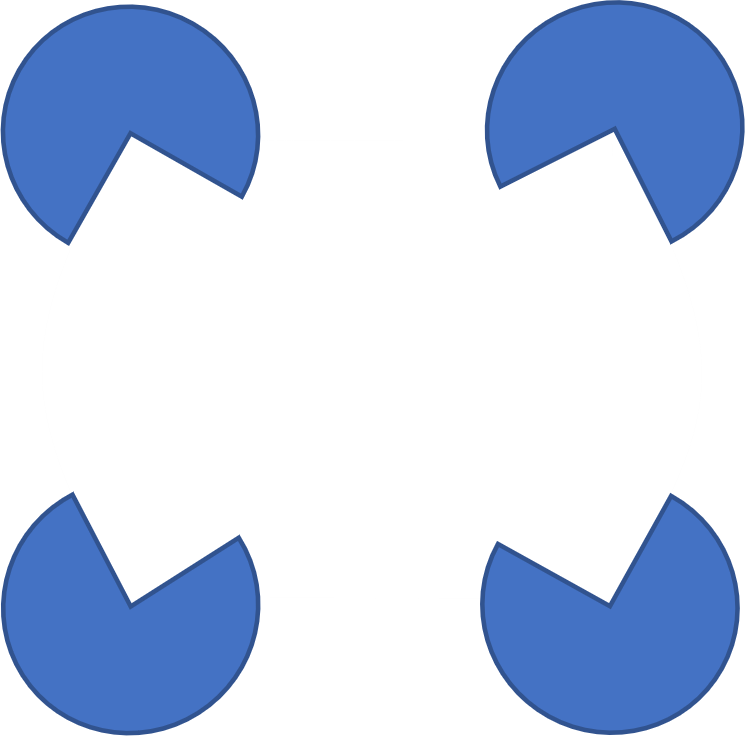3 | 4 | 1.1 cm / 1.1˚ | 90˚ | 4.2 cm (4.01°) ×2  3.3 cm (3.15°) ×2  5.5 cm (5.24°) (Feret) |
| 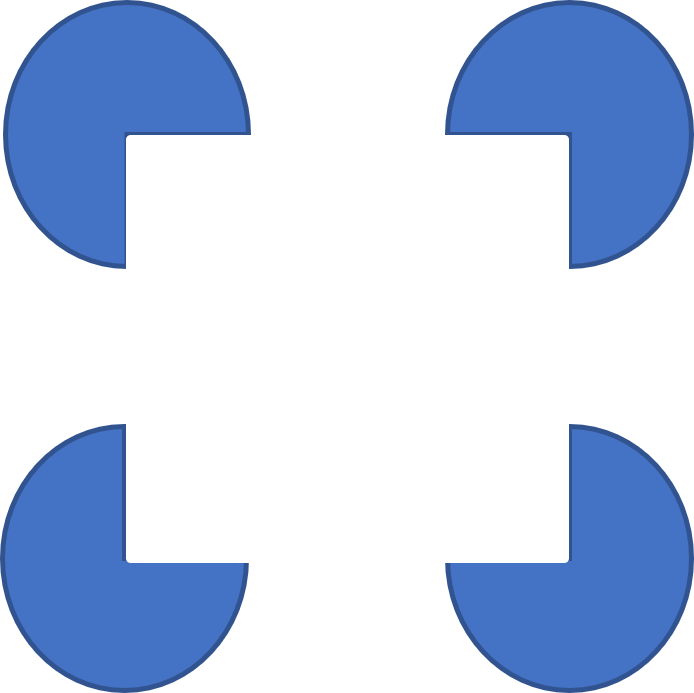4 | 4 | 1.1 cm / 1.1˚ | 90˚ | 4.2 cm (4.01°) ×2  3.7 cm (3.53°) ×2  5.0 cm (4.77°) (Feret) |
| 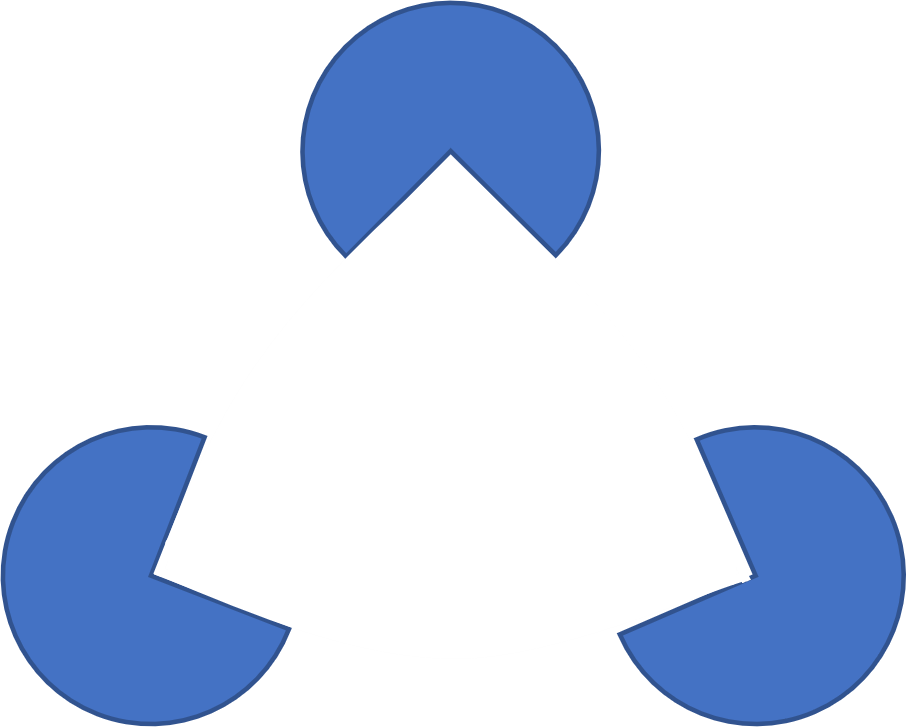5 | 3 | 1.0 cm / 1.0˚ | 90˚ | 4.0 cm (3.82°) ×2  4.4 cm (4.20°) ×1 |
| 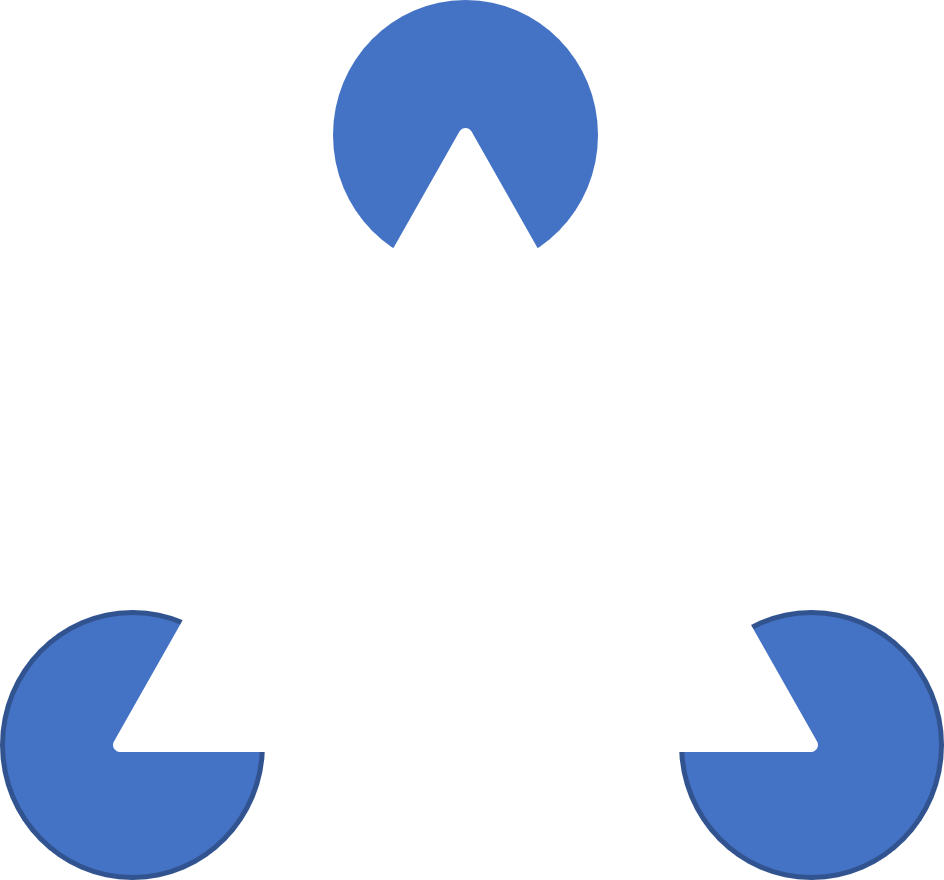6 | 3 | 1.0 cm /1.0˚ | 60˚ | 5.0 cm (4.77°) ×3 |
| 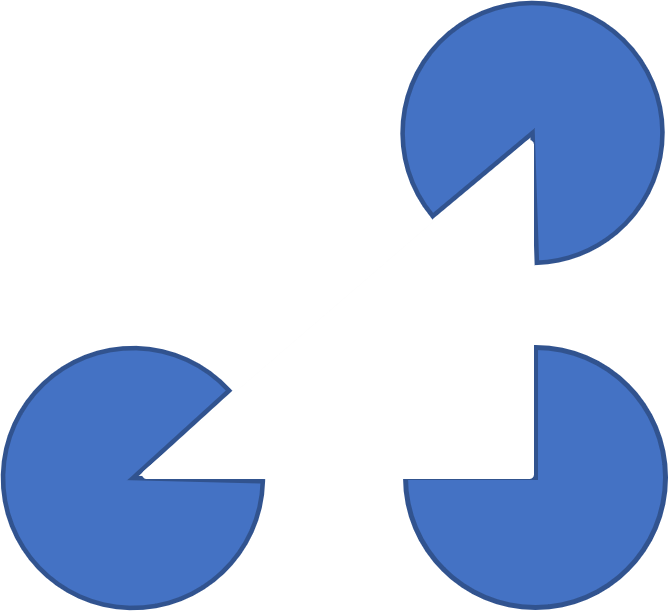7 | 3 | 1.1 cm / 1.1˚ | 90˚ and 45˚ | 4.0 cm (3.82°) ×1  3.9 cm (3.72°) ×1  5.2 cm (4.96°) (Feret) |
| 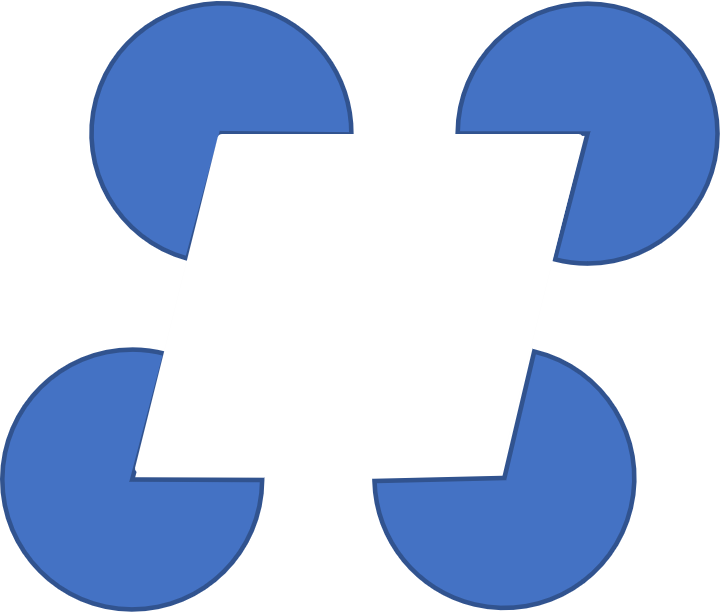8 | 4 | 1.1 cm / 1.1˚ | 77˚ and 103˚ | 3.4 cm (3.25°) ×4  5.3 cm (5.05°) (Feret) |
| 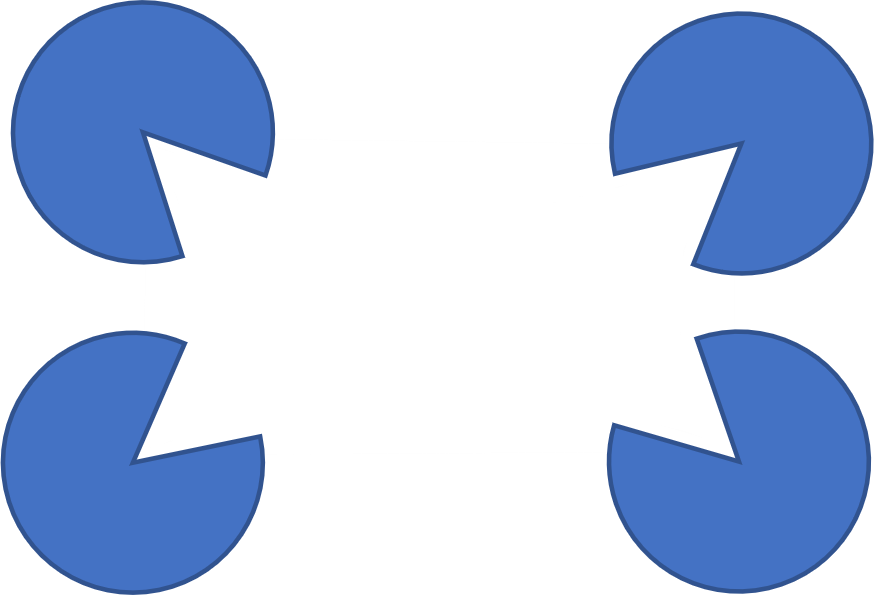9 | 4 | 1.1 cm / 1.1˚ | 55˚ | 4.5 cm (4.29°) ×2  3.0 cm (2.86°) ×2  5.5 cm (5.24°) (Feret) |

### Table S3c. Color Properties of Drag-and-Drop Stimuli

| Color Name | Stimulus Patch | Mean RGB (0–255) | HEX |
| --- | --- | --- | --- |
| Blue |  | (69, 114, 197) | #4572C5 |
| Green | 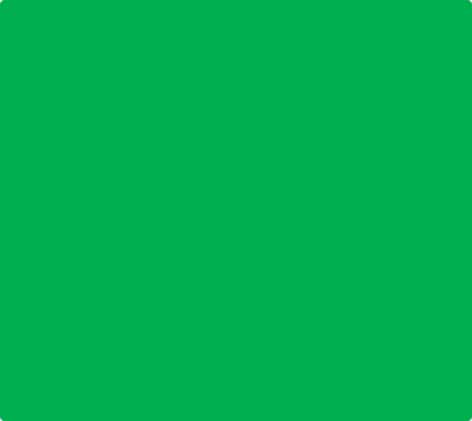 | (0, 175, 80) | #00AF50 |
| Yellow | 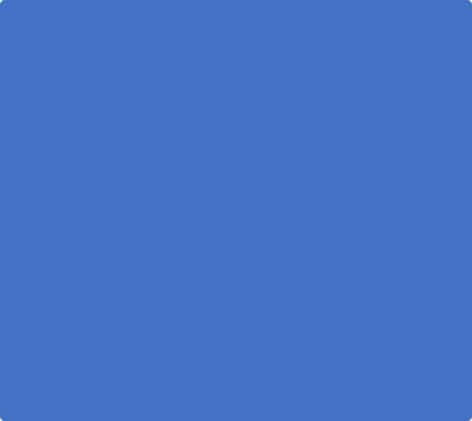 | (255, 255, 0) | #FFFF00 |
| Red |  | (254, 0, 2) | #FE0002 |
| Purple | 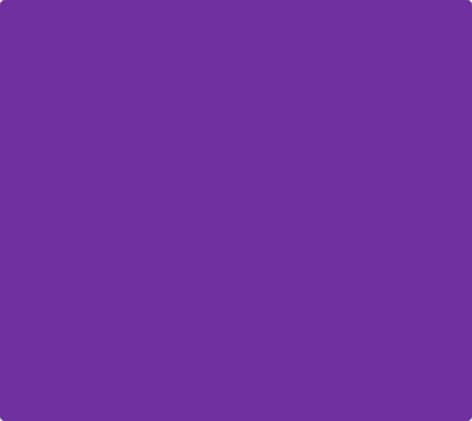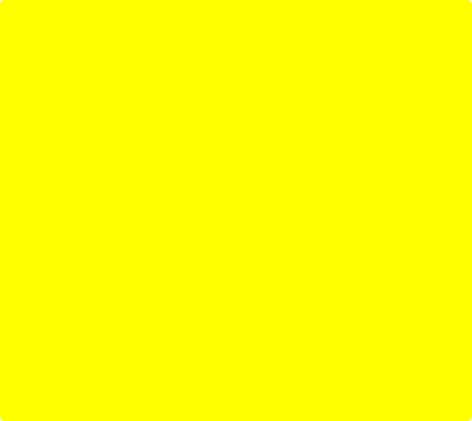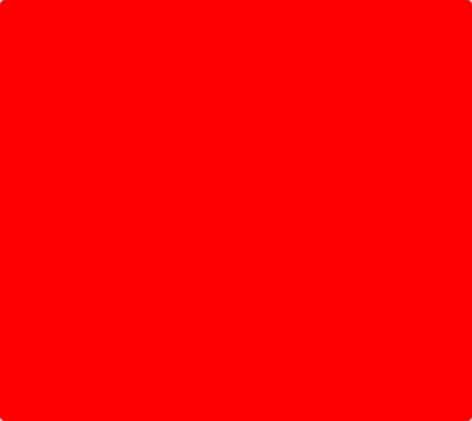 | (112, 48, 160) | #7030A0 |

Stimuli were displayed using fixed RGB values. No gamma correction or photometric calibration was applied, as the study did not involve luminance thresholds or contrast sensitivity measurements.
