## Supplemental Table S1 and S2 CC and KIC for "Minimally verbal children with autism may see the global, but point local: A behavioral and eye-tracking study in visual perceptual processing"

### Supplementary materials

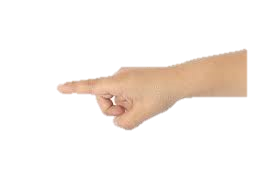

#### Table S1 Visual stimulus properties of Circular Contours (CCs)

| Properties | Contour-EASY | | Contour-MEDIUM | | Contour-HARD | |
| --- | --- | --- | --- | --- | --- | --- |
| Configurations | Low BG density* | | Medium BG density* | | High BG density* | |
| Dimensions | Diameter:4.3 cm(267px) | | Diameter:4.3 cm (267px) | | Diameter:4.3 cm (267px) | |
| Coordinates (x,y)   - peripheral L/R - Center (x) ↑/↓ | **P**:10.2 cm,2.75 cm  **C**: 0 cm, 2.75 cm | | **P:** 9.7 cm, 4.3 cm  **C**: 0 cm, 4.3 cm | | **P**: 9.7 cm, 4.3 cm  **C**: 0 cm, 4.3 cm | |
| Patch Contrast | Maximal | | Maximal | | Maximal | |
| Background | 60 cd/m² | | 60 cd/m² | | 60 cd/m² | |
| Visual angle   - Eccentricity - Size | **Preriheral**   - E:10° - S:4.1˚ | **Center (x)**   - E: 2.62˚ - S:4.1˚ | **Preriheral**   - E:10.1° - S:4.1˚ | **Center (x)**   - E: 4.1˚ - S: 4.1˚ | **Preriheral**   - E:10.1° - S:4.1˚ | **Center (x)**   - E: 4.1˚ - S: 4.1˚ |

*See text for details

#### Table S2 Visual stimulus properties of **Single Kanisza induced contours (KICs)**

| Stimulus Properties | | Inter-inducer center distance -size 1-3  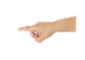Pointing | Coordinates of KICs size 1-3  Pointing | VAs of peripheral stimulus (peripheral):   - Eccentricity - Size | | Inter-inducer  center distance –  size 1-3  Eye-gaze  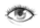 | Coordinates of KICs  size 1-3 | VAs of peripheral stimulus (peripheral):   - Eccentricity - Size | |
| --- | --- | --- | --- | --- | --- | --- | --- | --- | --- |
| ORIG |  | -1.8 cm  -2.9 cm  -4.2 cm | **S1:** 7.3 cm, 4.6 cm  **S2:** 6.5 cm, 6.3 cm  **S3:** 4.5 cm, 4.5 cm | **Peripheral**  S1:1.5˚  E1:8.2˚ | **Center**  S1:”-“  E1:0˚ | -2.9 cm  -4.2 cm  -6.5 cm | **S1:** 3.3 cm, 3.3 cm  **S2:** 4.5 cm, 4.5 cm  **S3:** 7.2 cm, 3.3 cm | **Peripheral**  S1:2.4˚  E1:4.5˚ | **Center**  S1:”-“  E1:0˚ |
| 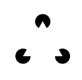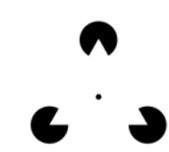  DOT |  |  |  | S2:2.4˚  E2:8.6˚ | S2:”-“  E2:0˚ |  |  | S2:3.5˚  E2:6.1˚ | S2:”-“  E2:0˚ |
| LUM |  |  |  | S3:3.5˚  E3:6.1˚ | S3:”-“  E3:0˚ |  |  | S3:5.4  E3:7.5 | S3: ”-“  E3: 0˚ |
| 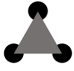  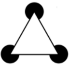  LINE | | Same | **S1:** 7.3 cm, 4.5 cm  **S2:** 5.7 cm, 5.0 cm  **S3:** 4.7 cm, 4.5 cm |  | | -2.8 cm  -3.8cm  -5.8 cm | **S1:** 3.3 cm, 3.0 cm  **S2:** 4.5 cm, 4.5 cm  **S3:** 7.1 cm, 3.0 cm | **Peripheral**  S1:2.3˚  E1:4.3˚ | **Center**  S1:”-“  E1:0˚ |
|  |  |  |  |  |  |  |  | S2:3.1˚  E2:6.1˚ | S2:”-“  E2:0˚ |
|  |  |  |  |  |  |  |  | S3:4.8  E3:7.8 | S3:”-“  E2:0˚ |
| Background luminance | | 335 cd/m² |  |  | | Same |  |  | |
| inducer | |  |  |  | |  |  |  | |

(Equilateral Kanizsa induced contour (KIC) triangles- 60˚ inducer opening)
