## Appendix1 Descriptive measures for "Minimally verbal children with autism may see the global, but point local: A behavioral and eye-tracking study in visual perceptual processing"

| **ID mvASD (pseudonyms)** | **AGE ^7^** | **Gender** | **Medication** | **L-VIS**  **Communication** | **SCQ** | **RCPM**  **Booklet^6^** | **RCPM**  **Puzzle^6^** | **shrieks** | **Non-verbal** | **Perseveration puzzle⁵** |
| --- | --- | --- | --- | --- | --- | --- | --- | --- | --- | --- |
| Lou | 8.75 | M | N | 7 | 26 | f.p. | f.p. | N | Y | 20 |
| Lars | 9.25 | M | Y | 9 | 18 | f.p. | 38.9 | Y | Y | 8 |
| Leif | 11.67 | M | N | 2 | 28 | f.p. | 13.8 | Y | Y | 4 |
| Len | 7.5 | M | N | 8² | 26 | f.p. | f.p. | Y | Y | 19 |
| Lee | 9.5 | M | Y | 5 | 24 | f.p. | f.p. | Y | Y | 15 |
| Lara | 8.3 | F | Y | 8 | 20 | f.p. | f.p. | Y | Y | 11 |
| Liam | 10.75 | M | ¹ | 6 | 23 | f.p. | f.p. | Y | Y | 8 |
| Lia | 8 | F | N | 5 | 26 | ³ | ³ | Y | Y | ³ |
| Leo | 10 | M | Y | 5 | 22 | f.p. | f.p. | Y | Y | 3 |
| Loni | 8.25 | M | Y | 6 | 30 | f.p. | f.p. | Y | N | 5 |
| Hal | 10 | M | N | 12 | 29 | 85 | 90 | N | N | 1 |
| Hila | 11.5 | F | Y | 14 | 18 | 28 | 34.4 | Y | N | 1 |
| Haya | 11.75 | F | N | 6 | 29 | 28 | 43.8 | Y | Y | 1 |
| Hugo | 12 | M | Y | 12 | 20 | f.p. | f.p. | Y | Y | 4 |
| Hank | 8.2 | M | N | 11 | 15 | 97 | 96.7 | N | Y | 0 |
| Hugh | 9.5 | M | Y | 13² | 22 | 95 | 97.5 | Y | N | 0 |
| Hana | 10,67 | F | Y | 8 | 23 | ³ | ³ | Y | Y | ³ |
| Hui | 11.3 | F | Y | 8 | 29 | f.p. | f.p. | Y | N | 12 |
| Hans | 8.5 | M | Y | 11 | 21 | f.p. | 7.5 | Y | N | 11 |
| Holm | 9.75 | M | Y | 11 | 27 | f.p. | 27.8 | N | Y | 2 |
| Hen | 7.5 | F | N | 12 | 25 | 44 | 75 | Y | Y | 2 |

### Table S4 Individual participants’ descriptive measures

¹Information not available. ²Scored by familiar staff. ³Unavailable for further testing. ⁴ Participant unwilling to complete testing. ⁵Perseveration: same answer location chosen on two consecutive items; RCPM never places the correct answer in the same location on adjacent items. f.p. = floor performance (less than 1^st^ percentile) = within age-range performance. ^6^ Percentile score in typical population (Ravens, 1998). ^7 Age at the time of testing KICs^

.
